# RyR2/IRBIT regulates insulin gene transcription, insulin content, and secretion in the insulinoma cell line INS-1

**DOI:** 10.1101/2021.11.24.469894

**Authors:** Kyle E. Harvey, Emily K. LaVigne, Mohd Saleem Dar, Amy E. Salyer, Evan P.S. Pratt, Paxton A. Sample, Uma Aryal, Humaira Gowher, Gregory H. Hockerman

## Abstract

The role of ER Ca^2+^ release via ryanodine receptors (RyR) in pancreatic β-cell function is not well defined. Deletion of RyR2 from the rat insulinoma INS-1 (RyR2^KO^) enhanced the Ca^2+^ integral (AUC) stimulated by 7.5 mM glucose, and rendered it sensitive to block by the IP_3_ receptor inhibitor xestospongin C, coincident with reduced levels of the protein IP_3_ <u>R</u>eceptor <u>B</u>inding protein released with Inositol 1,4,5 <u>T</u>risphosphate (IRBIT; aka AHCYL1). Deletion of IRBIT from INS-1 cells (IRBIT^KO^) increased the Ca^2+^ AUC in response to 7.5 mM glucose and induced xestospongin sensitivity. Insulin content and basal (2.5 mM glucose) and 7.5 mM glucose-stimulated insulin secretion were reduced in RyR2^KO^ cells and more modestly reduced in IRBIT^KO^ cells compared to controls. *INS2* mRNA levels were reduced in both RyR2^KO^ and IRBIT^KO^ cells, but *INS1* mRNA levels were specifically decreased in RyR2^KO^ cells. Nuclear localization of S-adenosylhomocysteinase (AHCY) was increased in RyR2^KO^ and IRBIT^KO^ cells. DNA methylation of the *INS1* and *INS2* gene promotor regions was very low, and not different among RyR2^KO^, IRBIT^KO^, and controls. In contrast, exon 2 of the *INS1* and *INS2* genes was more extensively methylated in RyR2^KO^ and IRBIT^KO^ cells than in controls. Proteomics analysis using LC-MS/MS revealed that deletion of RyR2 or IRBIT resulted in differential regulation of 314 and 137 proteins, respectively, with 41 in common. These results suggest that RyR2 regulates IRBIT levels and activity in INS-1 cells, and together maintain insulin content and secretion, and regulate the proteome, perhaps via DNA methylation.

**One sentence Summary:** Deletion of RyR2 from INS-1 cells had the unanticipated effect of reducing IRBIT proteins levels, and both RyR2 and IRBIT contribute to maintenance of glucose-stimulated insulin secretion.

## Introduction

Ca^2+^ signaling plays an essential role in pancreatic β-cell function and pathophysiology, and involves both Ca^2+^ influx into the cell via plasma membrane Ca^2+^ channels, and efflux of Ca^2+^ from the endoplasmic reticulum (ER) through ER membrane Ca^2+^ channels [1]. Voltage-gated Ca^2+^ channels Ca_v_1.2, Ca_v_1.3, and Ca_v_2.1 are located on the plasma membrane, and are key players in nutrient-stimulated insulin secretion [2] and in regulation of pancreatic β-cell proliferation [3]. The L-type channels Ca_v_1.2 and Ca_v_1.3 and the P/Q-Type channel Ca_v_2.1 are all expressed in human pancreatic β ls [2], but non-coding loss of function variants of CACNA1D, the gene that encodes Ca_v_1.3, are associated with an increased risk of type 2 diabetes [4].

The roles for the ER membrane Ca^2+^ channels ryanodine receptor 2 (RyR2) and inositol 1,4,5-triphosphate receptor (IP_3_R) are less well defined in β-cells. It was recently shown using targeted mass spectrometry that RyR2 is expressed in the INS-1 cell line and murine β-cells, and that ER stress affects RyR2 function [5]. Ca_v_1.2 activity is proposed to couple to RyR activation in INS-1 cells via Ca^2+^-induced Ca^2+^ release (CICR) [6]. In humans, mutations in RyR2 that cause Ca^2+^ “leak” induce both cardiac arrhythmias and glucose intolerance, and introduction of the corresponding mutations in mice led to glucose intolerance and a phenotype similar to type 2 diabetes [7]. A mutation in RyR2 mimicking CaMKII phosphorylation leads to basal hyperinsulinemia and glucose intolerance, both ascribed to increased leak of Ca^2+^ from the ER via RyR2 [8]. The major IP_3_R expressed in human β-cells is IP_3_R3 [9]. Expression of the ITPR3 gene is downregulated at the onset of type 1 diabetes [10], and animal studies support a role for IP_3_R3 in glucose-stimulated insulin secretion [11]. However, expression of IP_3_R1 and IP_3_R2, as well as IP_3_R3, were all elevated in human islets from type 2 diabetic donors, which was associated with increased ER stress, mitochondrial dysfunction and β-cell failure [12]. Both IP_3_R and RyR2 are Ca^2+^-activated ion channels that mediate release of Ca^2+^ of the ER, and regulate each other’s activity via Ca^2+^-dependent mechanisms [13].

Crosstalk between RyR2 and IP_3_R receptors may play a role in regulation of β-cell function [8]. One potential link in this crosstalk may be the regulation of the IP_3_ receptor binding protein IP_3_ <u>R</u>eceptor <u>B</u>inding protein released with Inositol 1,4,5 <u>T</u>risphosphate (IRBIT; a.k.a. AHCYL1). IRBIT binds to IP_3_ receptors in a manner competitive with IP_3_ and thus inhibits channel opening [14]. IRBIT must be serially phosphorylated to bind IP_3_ receptors, and the first step is a Ca^2+^-dependent phosphorylation thought to be mediated by a member of the Ca^2+^-calmodulin-dependent kinase family, possibly CaMKIV [15], but protein kinase D can also phosphorylate IRBIT *in vitro* [16]. Neither the mechanisms for activation of IRBIT *in vivo*, nor the role of IRBIT in pancreatic β-cells is known.

In the present study, we investigated the role of RyR2 in pancreatic β-cell signaling by deleting RyR2 from INS-1 cells using CRISPR-Cas9 gene editing [17]. We assessed the role of RyR2 in glucose stimulated Ca^2+^ transients, and found that RyR2 deletion increased the activity of IP_3_ receptors, co-incident with a marked decrease in protein levels of IRBIT. We further probed the role of IRBIT in several phenotypes of RyR2^KO^ cells by deleting IRBIT from INS-1 cells. Our results suggest that RyR2 regulates IRBIT activity, and together, they regulate insulin production and secretion.

## Results

### Characterization of RyR2^KO^ cells

Pancreatic β-cells potentially express RyR1, RyR2, and RyR3, but we chose to delete RyR2 in INS-1 cells given the previously reported effects of RyR2 mutations on glycemic control in both humans [7] and mice [8]. We used CRISPR-Cas9 gene editing, with two distinct guide RNAs, to introduce indels in exon 6 of the rat RyR2 gene in INS-1 cells that generated premature stop codons as assessed by sequencing genomic DNA (Fig 1A). Positive clones were identified by screening for loss of caffeine (5 mM) stimulation of RyR-mediated Ca^2+^ release from the ER in a 96-well format, using fura-2 AM (Fig1B). Single-cell Ca^2+^ measurements using fura-2 AM confirmed the loss of caffeine sensitivity in RyR2 knock out (RyR2^KO^) cells (Fig 1C); however, these cells displayed a strong increase in [Ca^2+^]_in_ in response to the muscarinic agonist carbachol (500 µM) (Fig 1C), suggesting that IP_3_ receptor activity is intact. Analysis of single-cell Ca^2+^ imaging experiments showed that the Ca^2+^ response (AUC) to caffeine was reduced by >90% in RyR2^KO^ cells, compared to control INS-1 cells (Fig 1D). The absence of RyR2 protein in RyR2^KO^ cells was confirmed by immunoblotting of microsomal proteins from control and RyR2^KO^ INS-1 cells and HEK 293 cells transfected with mouse RyR2-GFP [18] with a pan-specific RyR antibody (Fig 1E). Finally, basal (2.5 mM glucose) cytoplasmic Ca^2+^ levels as measured with fura-2 AM were slightly, but significantly lower in RyR2^KO^ cells compared to controls (Fig 1F), suggesting that RyR2 contributes to basal cytoplasmic Ca^2+^ levels. Thus, deletion of RyR2 in INS-1 cells abolishes the caffeine-stimulated Ca^2+^ response as well as the major species detected by immunoblotting with a pan-specific RyR antibody.

**Fig. 1.**
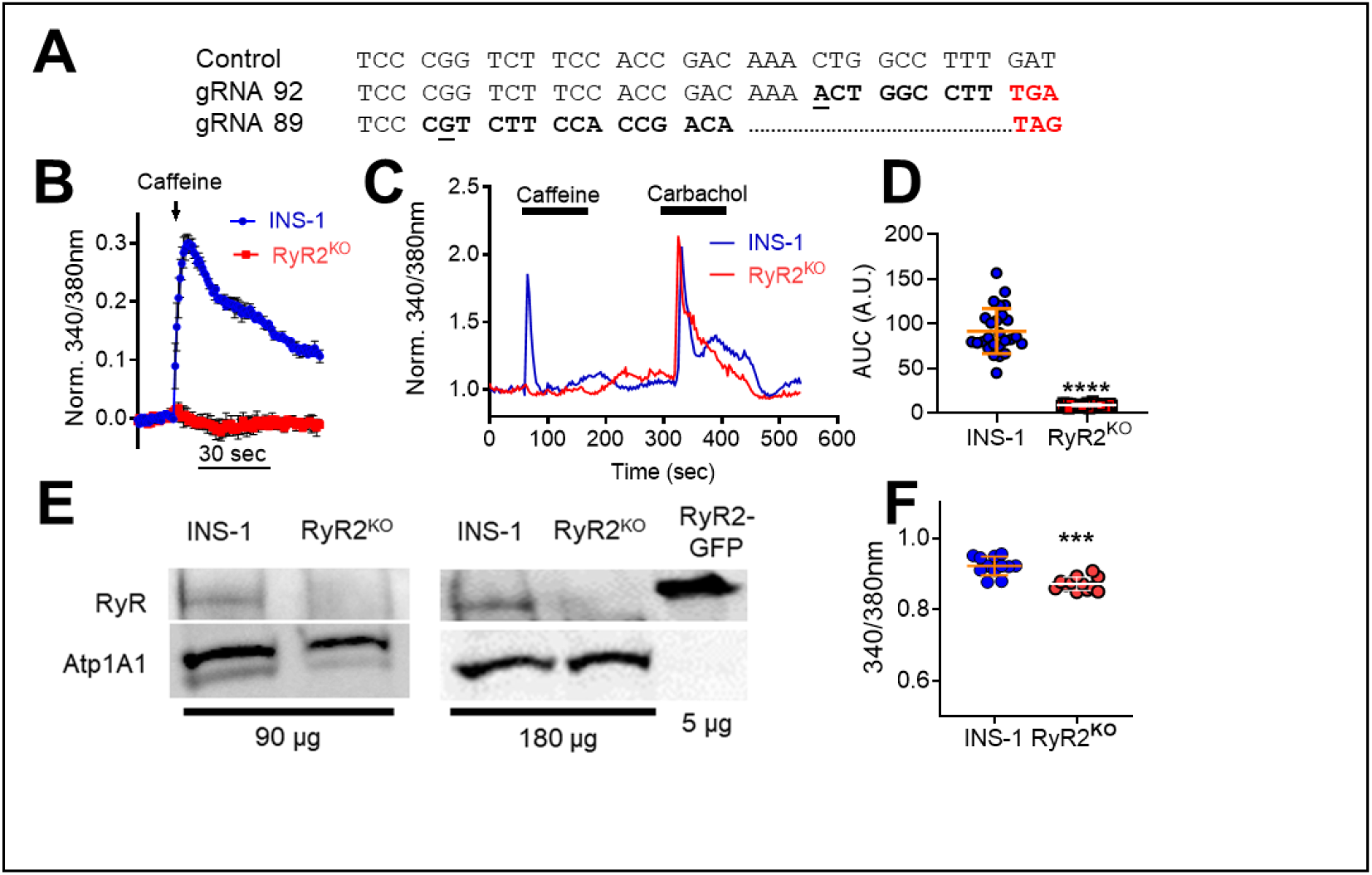
Characterization of RyR2^KO^ cells. **A)** Aligned genomic DNA sequences (exon 6) of the rat RYR2 gene in control and two distinct clones. Sequencing confirmed insertion of an indel (underlined/bolded) leading to a frameshift mutation (bolded) and a premature stop codon (red bolded). **B)** Preliminary screen (representative of 3 independent experiments) showing caffeine (5 mM) mobilization of Ca^2+^ in control INS-1 and an RyR2^KO^ clone measured with fura-2 AM using a 96-well plate format. Data are shown as mean ± SD. **C)** Representative experiments showing single-cell imaging of Ca^2+^ transients measured using fura-2 AM. RyR2^KO^ cells are insensitive to stimulation with the RyR2 agonist caffeine (5 mM), whereas caffeine elicits a rapid Ca^2+^ transient in INS-1 cells. Both cell lines display a robust Ca^2+^ transient in response to the muscarinic agonist carbachol (500 μM). **D)** Quantitation of the Ca^2+^ response (AUC) to 5 mM caffeine in control INS-1 and RyR2^KO^ cells. Lines represent mean ± SD ***, *P* < 0.0001, unpaired t-test; n = 26 cells (control), 30 cells (RyR2^KO^) from 3 independent experiments. **E)** Immunoblots for RyR from control and RyR2^KO^ cells. The total protein loaded in each lane (μg) is indicated. Samples were probed with a pan-specific RyR antibody. As a positive control, recombinant mouse RyR2 fused to GFP and expressed in HEK 293T cells was included. As a loading control, membranes were also probed with an antibody to the α1 subunit of the Na+/K+- Atpase (Atp1A1). Immunoblots shown are representative of 4 independent experiments. **F)** Basal Ca^2+^ levels in control INS-1 and RyR2^KO^ cells measured with fura-2 AM. ***, *P* < 0.001; unpaired t-test, n = 12 (control and RyR2^KO^) over 3 independent experiments. Lines represent mean ± SD.

### Ca^2+^ dynamics in RyR2^KO^ cells

Stimulation of both control INS-1 and RyR2^KO^ cells with 7.5 mM glucose resulted in periodic [Ca^2+^]_in_ oscillations (Fig 2A). The Ca^2+^ integral was inhibited in both cell lines by 2 µM nicardipine; however, the total AUC in the absence of nicardipine was greater in RyR2^KO^ cells compared to control cells (Fig 2B). The Ca^2+^ response to 7.5 mM glucose was inhibited by 1 µM xestospongin C in RyR2^KO^ cells but not in controls (Fig 2C, D & E). Xestospongin C also increased the time between peaks in RyR2^KO^ cells (Fig 2F), but reduced the time between peaks in control INS-1 cells (Fig 2F). Thus, IP_3_ receptors are major contributors to glucose-stimulated Ca^2+^ oscillations in RyR2^KO^ cells, but not in control INS-1 cells. However, IP_3_ receptors regulate the timing of oscillations in response to glucose in both control and RyR2^KO^ cells, but with opposite effect.

**Fig. 2.**
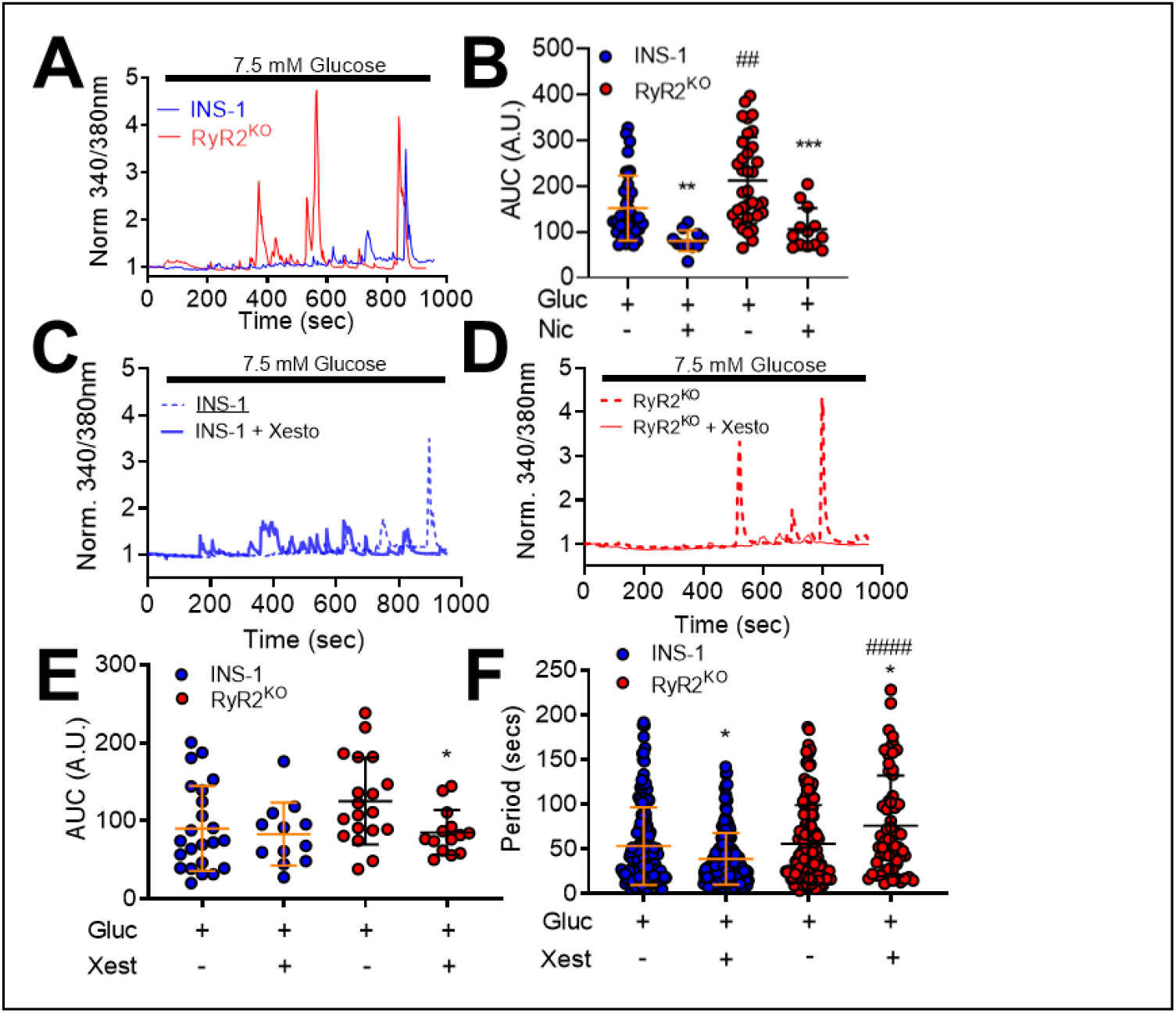
Glucose-stimulated Ca^2+^ transients in INS-1 and RyR2^KO^ cells. **A)** Single-cell Ca^2+^ transients measured in INS-1 and RyR2^KO^ cells in response to 7.5 mM glucose. **B)** Nicardipine (Nic; 2 μM) inhibited the Ca^2+^ AUC in INS-1 and RyR2^KO^ INS-1 cells stimulated with 7.5 mM glucose. Additionally, glucose-stimulated Ca^2+^ AUC was increased in RyR2^KO^ INS-1 cells as compared to INS-1 cells. INS-1: glucose, n = 35; glucose + Nicardipine, n = 11. RyR2^KO^: glucose, n = 34; glucose + nicardipine, n = 13 from 3 independent experiments (**, *P =* 0.0022; ***, *P* = 0.0003; Students unpaired t-test glucose alone compared to glucose + nicardipine. ^##^, *P* = 0.0059; one-way ANOVA with Tukey’s *post-hoc* test) INS-1 + glucose compared to RyR2^KO^ + glucose. Lines represent mean ± SD. Representative single-cell Ca^2+^ transients measured in INS-1 cells **(C)** and RyR2^KO^ INS-1 cells **(D)** stimulated with glucose in the presence or absence of xestospongin C (Xesto; 1 μM). **E)** Xestospongin C (Xest) significantly diminished the glucose-stimulated Ca^2+^ AUC in RyR2^KO^ INS-1 cells but had no effect in INS-1 cells (*, *P* < 0.0426) (two-way ANOVA with Sidak’s *post-hoc* test). INS-1: glucose, n = 23 cells; glucose + xest, n = 12 cells in 5 and 3 independent experiments, respectively. RyR2^KO^: glucose, n = 19 cells; glucose + xest, n = 14 cells in 5 and 3 independent experiments, respectively. Lines represent mean ± SD. **F)** The presence of xestospongin c reduced the time between glucose- stimulated Ca^2+^ oscillations (period) in INS-1 cells, while it significantly increased the period in RyR2^KO^ INS-1 cells. Furthermore, xestospongin c increased the period in RyR2^KO^ INS-1 cells as compared to INS-1 cells (^####^, *P* < 0.0001, *, *P* = 0.011 (INS-1), *, *P* = 0.015 (RyR2^KO^)) (two-way ANOVA with Tukey’s *post-hoc* test). INS-1: glucose, n = 159 peaks; glucose + xest, n = 153 peaks in 5 independent experiments. RyR2^KO^: glucose, n = 160 peaks; glucose + xest, n = 62 peaks in 5 independent experiments). Lines represent mean ± SD.

### Regulation of IRBIT by RyR2

Given the apparent increase in IP_3_ receptor activation in response to glucose that we observed in RyR2^KO^ cells, we examined the ability of glucose to activate phospholipase C (PLC) in both RyR2^KO^ and INS-1 cells. Stimulation with 7.5 mM glucose (Fig 3A) or 500 µM the muscarinic receptor agonist carbachol (Fig 3B) stimulated PLC activity above basal levels in both control and RyR2^KO^ cells. However, stimulated PLC activity was decreased in RyR2^KO^ cells compared to control INS-1 cells (Fig 3C). Thus, the increased IP_3_ receptor activity observed with glucose stimulation in RyR2^KO^ cells is unlikely to result from increased PLC activity and greater accumulation of IP_3_. Total cellular phosphatidylinositol bisphosphate (PIP_2_) levels in fixed, saponin-treated cells was measured using immunocytochemistry. RyR2^KO^ cells contained slightly more PIP_2_ than control INS-1 cells (Fig 3D). Thus, the reduced PLC activity in RyR2^KO^ cells is unlikely to be the result of limiting substrate levels. Given these findings, we measured the levels of the protein IRBIT (aka AHCYL1). IRBIT binds to IP_3_Rs and competitively inhibits IP_3_ binding [14]. Moreover, IRBIT must be phosphorylated by a Ca^2+^-dependent kinase in order to bind IP_3_Rs [15]. Using semi-quantitative immunoblotting, we found that IRBIT protein levels normalized to actin, were substantially reduced (∼80% reduction) in RyR2^KO^ cells compared to control cells (Fig 3E). These data suggest that RyR2 plays a role in regulating IRBIT activity, such that, in the absence of RyR2, IRBIT protein levels/activity are suppressed, allowing hyperactivation of IP_3_Rs.

**Fig. 3.**
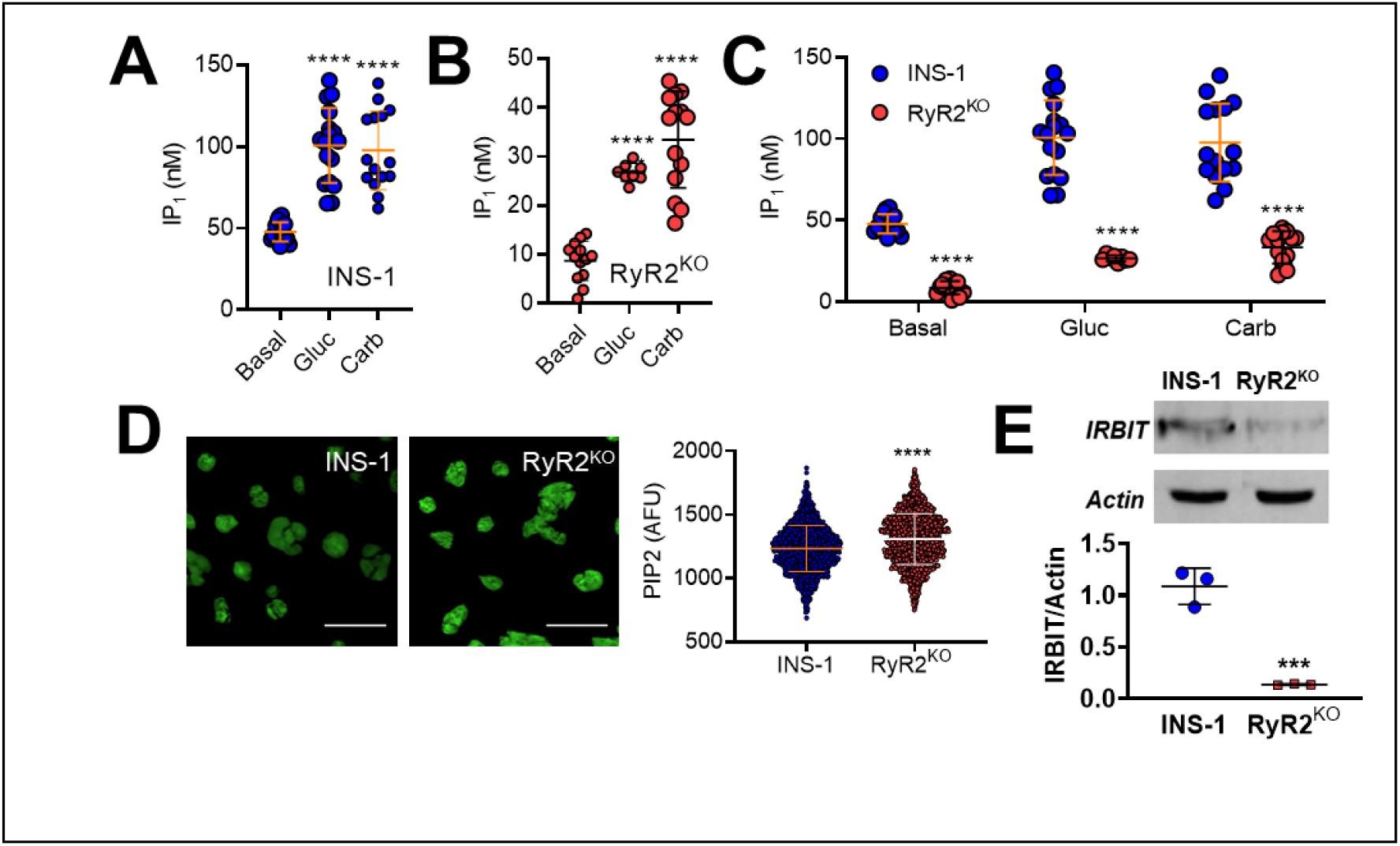
Phospholipase C activity in control INS-1 and RyR2^KO^ cells-. PLC activity was assessed by measuring basal and stimulated IP_1_ levels (an indirect measure of IP_3_ levels) in **A)** INS-1 and **B)** RyR2^KO^ INS-1 cells. 1 hour stimulation with 7.5 mM glucose (Gluc) or 500 μM carbachol (Carb;) resulted in increased IP_1_ accumulation in control INS-1 cells compared to basal (A; ****, *P* < 0.0001; One-way ANOVA, Dunnett’s *post-hoc* test (basal, n = 16; Gluc, n = 17; Carb, n = 15 in 4 independent experiments done in triplicate) and RyR2^KO^ cells (B; *, *P* = 0.0107, ****, *P* < 0.0001; One-way ANOVA, Dunnett’s *post-hoc* test; (basal, n = 12; Gluc, n = 8; Carb, n = 14 in 3 independent experiments done in triplicate). Lines represent means ± SD. **C)** Basal and stimulated IP_1_ accumulation was significantly reduced in RyR2^KO^ INS-1 cells as compared to INS-1 cells (****, *P* < 0.0001; Two-way ANOVA, Šídák’s *post-hoc* test). Data re- plotted from **A** and **B** for comparison. **D)** Representative images of fixed, saponin permeabilized control INS-1 and RyR2^KO^ cells stained with a primary antibody against PIP_2_, followed by IgG-κ binding protein conjugated to CFL 488. Scale bars = 50 μm. The mean fluorescence intensity was slightly greater in RyR2^KO^ cells compared to control INS-1 cells. ****, *P* < 0.0001 (unpaired t-test) INS-1: n = 1294 cells; RyR2^KO^: n = 1377 cells from 3 independent experiments. Lines represent mean ± SD. **E)** Representative immunoblots of IRBIT in control INS-1 and RyR2^KO^ cells (top) and quantitation of IRBIT levels, normalized to actin, in control INS-1 cells and RyR2^KO^ cells (n = 3). ***, *P* < 0.0007 (unpaired t-test). Lines represent mean ± SD.

### Characterization of IRBIT^KO^ cells

To decipher which effects of RyR2 deletion are likely directly due to loss of ER Ca^2+^ release via RyR2, and which are likely mediated by dysregulation of IRBIT, we deleted IRBIT from INS-1 cells using CRISPR/cas9 gene editing with gRNAs targeted to exon 6 of the AHCYL1 gene.

Genomic DNA sequencing identified clones with expected indels (Fig 4A). Immunoblots of cell lysates from IRBIT^KO^ cells confirmed the absence of IRBIT protein (Fig 4B). IRBIT mRNA was reduced in IRBIT^KO^ cells compared to controls, but IRBIT mRNA levels were not reduced in RyR2^KO^ cells (Fig 4C). Basal (2.5 mM glucose) [Ca^2+^]_in_ measured with fura-2 AM was greater in IRBIT^KO^ cells compared to RyR2^KO^ cells, but was not different from that measured in control INS-1 cells (Fig 4D). The [Ca^2+^]_in_ response to 5 mM caffeine in IRBIT^KO^ cells, measured with fura-2 AM, was reduced compared to control INS-1 cells, but was much greater than that observed in RyR2^KO^ cells (Fig 4E). Using the ER-targeted Ca^2+^ indicator D1ER [19] to measure ER Ca^2+^ levels, we found that basal ER [Ca^2+^] was reduced in IRBIT^KO^ cells compared to both RyR2^KO^ and control INS-1 cells, and that thapsigargin treatment reduced ER [Ca^2+^] to levels that were not different across the three cell lines (Fig 4F). The [Ca^2+^]_in_ response to 7.5 mM glucose in IRBIT^KO^ cells was inhibited by 1 µM xestospongin C (Fig 4G & 4H), and was greater than in control INS-1 cells, but not different from that measured in RyR2^KO^ cells (Fig 4I). Thus, the near abolition of the caffeine response in RyR2^KO^ cells is the direct result of the deletion of RyR2, but IRBIT is required to maintain the full magnitude of RyR2-mediated Ca^2+^ release. In contrast, increased Ca^2+^ response to 7.5 mM glucose and block of this response by xestospongin C in RyR2^KO^ cells is likely the result of reduced IRBIT levels.

**Fig 4.**
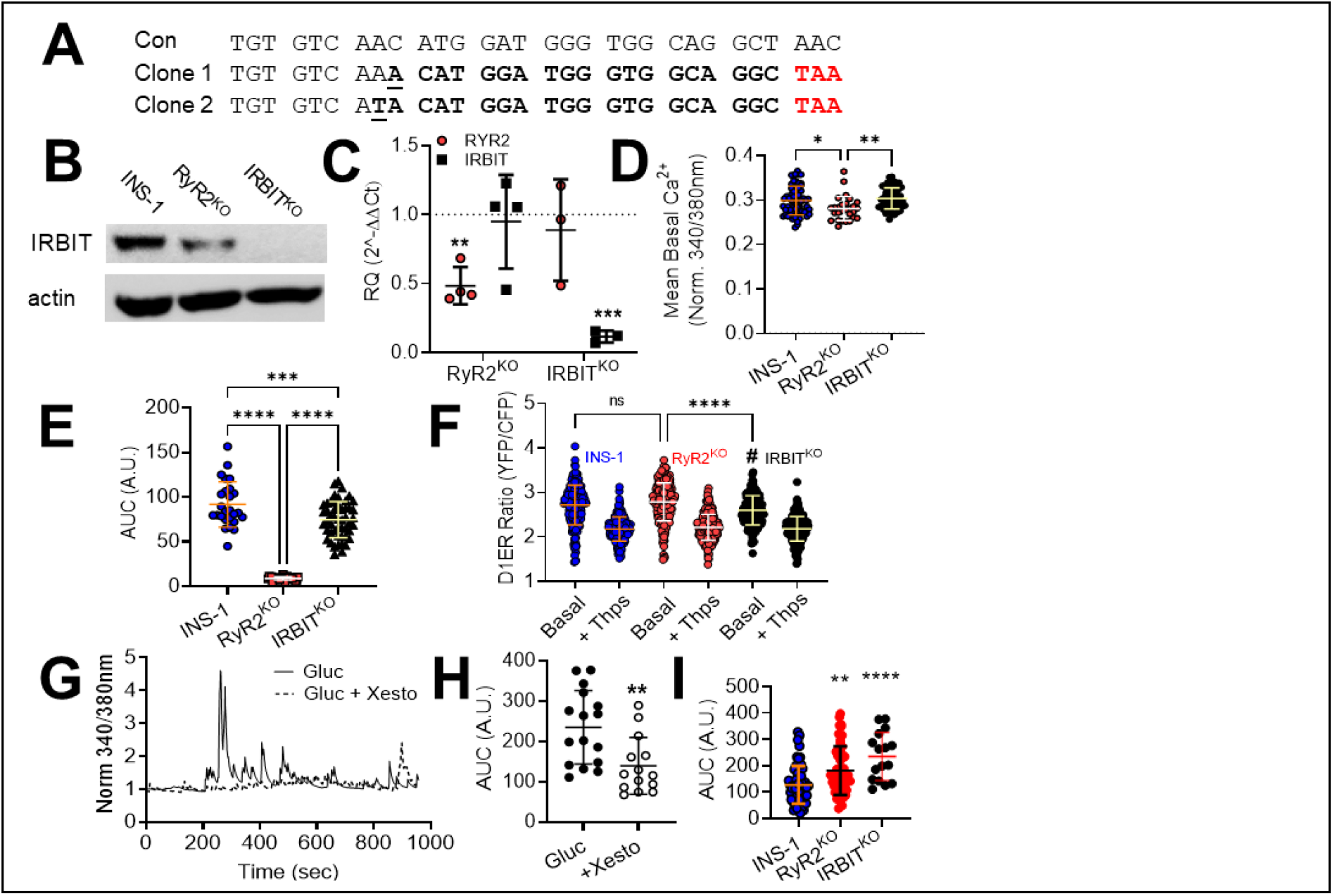
Characterization of IRBIT^KO^ cells-. **A)** Genomic sequence within exon 6 of the rat AHCYL1 gene with sequences from two clones derived from CRISPR-Cas9 gene editing of INS-1 cells. The underlined nucleotides indicate the site of the indel induced by gene editing in each clone. Each indel induced a frameshift resulting in the premature stop codon shown in red. **B)** Immunoblot for IRBIT in control INS-1 cells, RyR2^KO^ cells, and IRBIT^KO^ cells. Blot shown is representative of 4 independent experiments. **C)** RyR2 and IRBIT mRNA levels in RyR2^KO^ and IRBIT ^KO^ cells relative to control INS-1 cells. Each point represents an independent experiment done in triplicate. RyR2^KO^, n = 4, IRBIT^KO^ n = 3 (*\*\*\*, P* = 0.0008, **, P = 0.0046, One-sample t-test). **D)** Basal levels of cytoplasmic Ca^2+^ as assessed with fura-2 AM (340/380 nm). INS-1: n = 67 cells; RyR2^KO^: n = 28 cells ; IRBIT^KO^: n = 60 cells from 3 independent experiments. **, *P* = 0.002; *, *P* = 0.015 (one-way ANOVA with Tukey’s *post-hoc* test). **E)** Ca^2+^ release stimulated by 5 mM caffeine (AUC) in control INS-1, RyR2^KO^, and IRBIT^KO^ cells. n = 26 cells (control), 30 cells (RyR2^KO^), 50 cells (IRBIT^KO^). ****, *P* < 0.0001; ***, *P* = 0.0008 (one-way ANOVA with Tukey’s *post-hoc* test). Control and RyR2^KO^ data are the same as presented in Figure 1D. **F)** ER Ca^2+^ levels measured with D1ER (ratio of YFP:CFP fluorescence intensity) before or after the addition of 1 µM thapsigargin. ****, *P* < 0.0001; *, *P* = 0.019. INS-1: basal n = 217 cells, thaps n = 230 cells ; RyR2^KO^: basal n = 229 cells, thaps n = 296 cells; IRBIT^KO^: basal n = 147 cells, thaps n = 272 cells from 3 independent experiments. (one-way ANOVA with Tukey’s *post-hoc* test). There was no significant difference in ratios between thapsigargin-treated INS-1, RyR2^KO^, or IRBIT^KO^ cells (one-way ANOVA). **G)** Representative time courses of Ca^2+^ oscillations stimulated by 7.5 mM glucose in IRBIT^KO^ cells in the presence or absence of 1 µM xestospongin C. **H)** Xestospongin C reduces the Ca^2+^ response (AUC) to 7.5 mM glucose in IRBIT^KO^ cells. **, *P* = 0.0037; Glucose n = 16 cells, Glucose + Xesto n = 14 cells from 3 independent experiments (unpaired t-test). **I)** Comparison of Ca^2+^ integral (AUC) upon stimulation with 7.5 mM glucose in control INS-1, RyR2^KO^, and IRBIT^KO^ cells. ****, *P* < 0.0001; **, *P* = 0.0024 (one-way ANOVA with Tukey’s *post-hoc* test). INS-1 n = 58 cells, RyR2^KO^ n = 53 cells, IRBIT^KO^ n = 16 cells from 3 independent experiments, respectively. Data from INS-1 and RyR2^KO^ cells shown in Figs. 2B and 2E, were combined and replotted with data from IRBIT^KO^ cells for comparison. Lines represent mean ± SD.

### Insulin content and secretion in RyR2^KO^ and IRBIT^KO^ cells

Since [Ca^2+^]_in_ is a key regulator of insulin secretion, we measured insulin secretion in response to glucose in control INS-1, RyR2^KO^, and IRBIT^KO^ cells. We examined glucose-stimulated insulin secretion (GSIS) at 2.5 mM and 7.5 mM glucose in all three cell lines. Given the large contribution of IP_3_ receptors to the Ca^2+^ response to glucose in RyR2^KO^ and IRBIT^KO^ cells, we examined the contribution of L-type Ca^2+^ channels and IP_3_ receptors to GSIS. We found that as in control cells, 2 µM nicardipine (L-type channel blocker) completely inhibited 7.5 mM GSIS in all three cell lines, but that 1 µM xestospongin C (IP_3_ receptor antagonist) did not affect 7.5 mM GSIS in any of the cell lines. In each case, nicardipine suppressed GSIS to a level not different from that stimulated by 2.5 mM glucose (Fig 5A). Insulin secretion at both 2.5 mM and 7.5 mM glucose was reduced in both RyR2^KO^ and IRBIT^KO^ cells compared to controls (Fig 5B). Insulin content was substantially reduced (∼70% decrease) in RyR2^KO^ cells and more modestly reduced (∼40% decrease) in IRBIT^KO^ cells compared to control INS-1 cells as measured by insulin assay of ethanol/HCl-extracted cells, normalized to protein (Fig 5C). ER Ca^2+^ release contributes to glucose-dependent activation of extracellular-signal regulated protein kinase (ERK) 1/2 [3], which phosphorylates and activates several transcription factors involved in positive regulation of insulin transcription [20]. Therefore, we asked if glucose-stimulated ERK 1/2 phosphorylation was disrupted in RyR2^KO^ or IRBIT^KO^ cells (Fig 5D). 7.5 mM glucose stimulated an ∼3-fold increase in pERK 1/2 compared to 2.5 mM glucose in control, RyR2^KO^, IRBIT^KO^ cells (Fig. 5D). Epidermal growth factor (15 nM) potentiated pERK in the presence of 7.5 mM glucose in control and RyR2^KO^ cells, but this potentiation was abolished in IRBIT^KO^ cells (Fig 5D). Thus, deficits in glucose-stimulated ERK 1/2 phosphorylation likely don’t contribute to the decreased insulin content in RyR2^KO^ or IRBIT^KO^ cells. However, levels of *INS2* transcript were markedly suppressed in both RyR2^KO^ and IRBIT^KO^ cells compared to controls (Fig 5E). In contrast, the levels of *INS1* mRNA were not different from controls in IRBIT^KO^ cells, but were decreased in RyR2^KO^ cells compared to controls (Fig 5E).

**Fig. 5.**
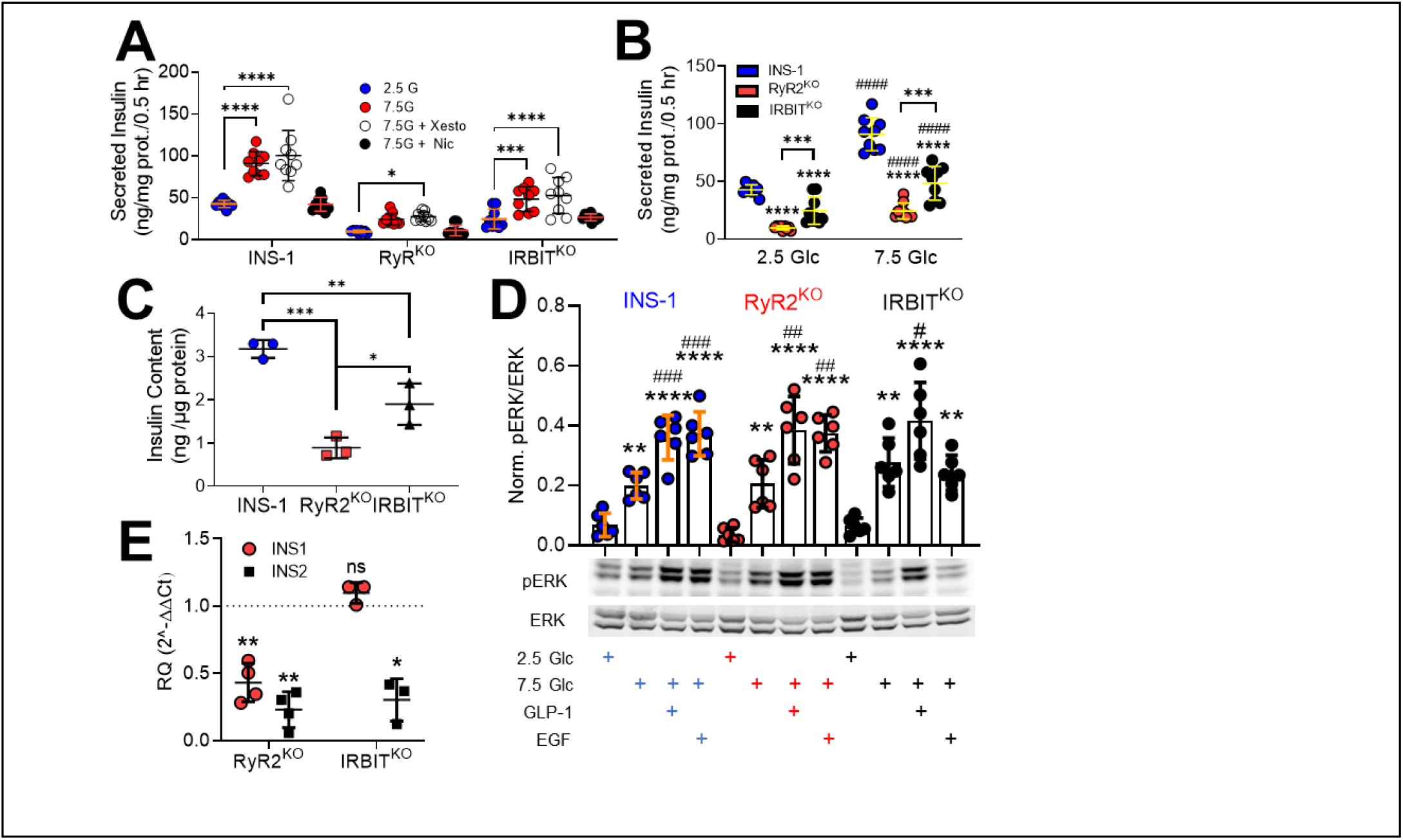
RyR2/IRBIT regulate insulin secretion and content. **A)** Insulin secretion stimulated by 7.5 mM glucose in INS-1, RyR2^KO^, and IRBIT^KO^ cells is inhibited by 2 µM nicardipine, but not 1 µM xestospongin C. ****, *P* < 0.0001; ***, *P* = 0.0008; *\*, P* = 0.017 (n = 9 from 3 independent experiments). (two-way ANOVA with Dunnett’s post-hoc test.) **B)** Insulin secretion at both 2.5 mM and 7.5 mM glucose were reduced by RyR2 deletion and IRBIT deletion. ^####^, *P* < 0.0001 compared to 2.5 mM glucose; ****, *P* < 0.0001; ***, *P* < 0.001 compared to INS-1 (n = 9 from 3 independent experiments). (two-way ANOVA with Tukey’s *post-hoc* test). **C)** Insulin content, normalized to total cellular protein in INS-1, RyR2^KO^, and IRBIT^KO^ cells. ***, *P* = 0.0004; **, *P* = 0.0076; *, *P* = 0.0221 (n = 3) (one-way ANOVA with Tukey’s *post-hoc* test). **D)** Phosphorylation of ERK1/2 in response to 2.5 mM or 7.5 mM glucose (Glc), 7.5 mM glucose + 50 nM GLP-1, or 7.5 mM glucose + 15 nM EGF. Immunoblot shown is representative of 6 independent experiments. Quantitation- INS-1: **, *P =* 0.0058; ****, *P* < 0.0001 compared to 2.5 mM glucose; ^###^, *P* = 0.0008 (GLP-1 + 7.5 mM Glc) and *P* = 0.0004 (EGF + 7.5mM Glc) compared to 7.5 mM glucose. RyR2^KO^: **, *P =* 0.0048; ****, *P* < 0.0001 compared to 2.5 mM glucose; ^##^, *P* = 0.0031 (GLP-1 + 7.5 mM Glc) and *P* = 0.0056 (EGF + 7.5 mM Glc) compared to 7.5 mM glucose. IRBIT^KO^: **, *P =* 0.0012; ****, *P* < 0.0001, **, *P* = 0.0064 compared to 2.5 mM glucose; ^#^, *P* = 0.0383 (GLP-1 + 7.5 mM Glc) compared to 7.5 mM glucose (one-way ANOVA with Tukey’s *post-hoc* test, n = 6 separate experiments for each group). **E)** mRNA levels of *INS1* and *INS2* transcript in RyR2^KO^ and IRBIT^KO^ cells relative to control INS-1 cells, as assessed by rt-qPCR. RyR2^KO^: INS1, **, *P* = 0.0042 (n = 4); INS2, **, *P* = 0.0014 (n = 4). IRBIT^KO^: INS2, *, *P* = 0.0164 (n = 3). (one-sample t-test). Glyceraldehyde-3-phosphate dehydrogenase mRNA was used as the reference for the data shown. Equivalent results were obtained using phosphoglycerate kinase mRNA as the reference (data not shown). Lines represent mean ± SD.

### Regulation of AHCY localization by RyR2 and IRBIT

One potentially global effect of IRBIT deletion is dysregulation of methyltransferase activity. IRBIT binds to S-adenosyl homocysteinase (AHCY) [21, 22], the only enzyme known to hydrolyze S-adenosyl homocysteine (SAH) and relieve product inhibition of DNA, RNA, and protein methyltransferases, and is reported to regulate nuclear localization of AHCY [22]. To determine if IRBIT thus regulates AHCY in INS-1 cells, we examined the subcellular localization of AHCY in control, RyR2^KO^, and IRBIT^KO^ cells using immunocytochemistry. Confocal micrographs were taken of fixed cells labeled with a primary AHCY antibody, and an Alex Fluor 488-conjugated secondary antibody, counterstained with Hoechst 33342 to define the nuclei (Fig 6A). Control experiments in which primary antibodies were omitted resulted in cells with negligible Alex Fluor 488 fluorescence (Fig 6B). In control INS-1 cells, AHCY is preferentially localized in the nucleus relative to the cytoplasm (4:1 ratio) (Fig 6A&C). In RyR2^KO^ cells, AHCY localized to the nucleus relative to the cytoplasm is (4:1 ratio), but more AHCY was detected in the nucleus compared to control INS-1 cells. Deletion of IRBIT resulted in a marked depletion of AHCY in the cytoplasm, resulting in a nucleus to cytoplasm ratio of 13:1 (Fig 6A&C). However, the total amount of AHCY detected in the nucleus of IRBIT^KO^ cells was reduced compared to control INS-1 cells (Fig 6C). Thus, deletion of RyR2 or IRBIT correlates with either increased nuclear AHCY (RyR2^KO^ cells), or a sharp decrease in non-nuclear localized AHCY (IRBIT^KO^ cells). Increased accumulation of AHCY in the nucleus could potentially increase DNA methyltransferase activity (Fig 6D).

**Fig. 6.**
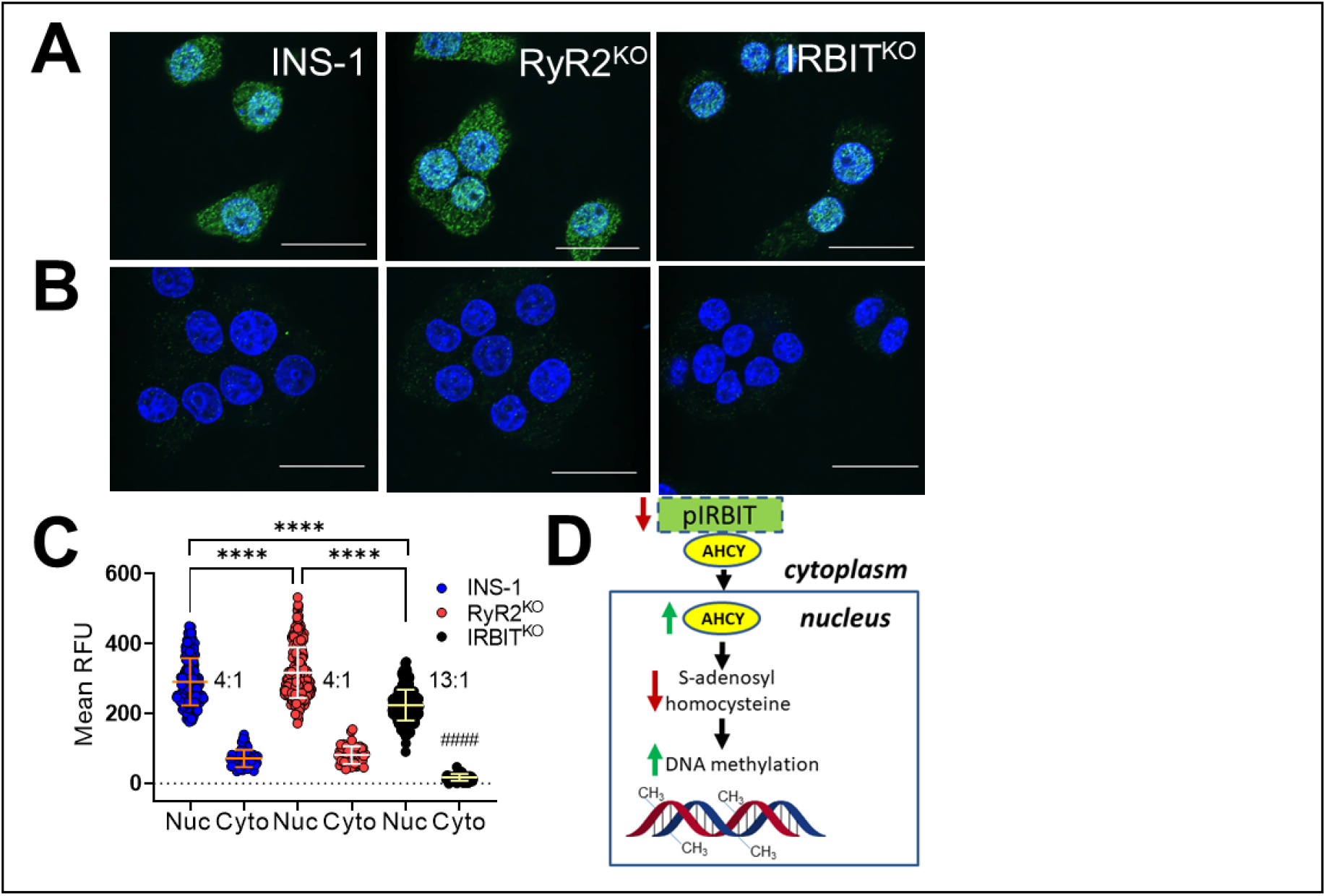
RyR2/IRBIT regulate nuclear localization of AHCY. **A)** Images of indicated cells fixed and stained with anti-AHCY antibodies and Alexa Fluor 488-conjugated secondary antibodies. Nuclei are stained with Hoechst 33342. **B)** Images of indicated cells fixed and stained as in A, except that the primary antibodies were omitted. Scale bars = 20 μm. **C)** Quantification of Alexa Fluor 488 fluorescence intensity in nuclei vs outside of nuclei in images of cells stained as in A. n = 280 cells for INS-1 nucleus; n = 280 cells for RyR2^KO^ nucleus; n = 373 for IRBIT^KO^ nucleus; n = 50 for INS-1 cytoplasm; n = 55 cells for RyR2^KO^ cytoplasm; n = 53 cells for IRBIT^KO^ cytoplasm from 3 independent experiments. Inset numbers represent the nucleus:cytoplasm ratio of Alexa Fluor 488 fluorescence intensity. ****, *P* < 0.0001; ^####^, *P* < 0.0001 compared to both INS-1 cyto and IRBIT cyto (one-way ANOVA with Tukey’s *post-hoc* test). Lines represent mean ± SD. **D)** Model for the regulation of DNA methylation by loss of IRBIT.

### Regulation of *INS1* and *INS2* gene methylation by RyR2 and IRBIT

Given the changes in nuclear localization of AHCY in RyR2^KO^ and IRBIT^KO^ cells, we examined the possibility that insulin genes in the knockout cells were differentially methylated. PCR amplification of genomic DNA regions, with or without digestion by a methylation-dependent endonuclease using primers that flank potential methylation sites (Fig 7A&B), provide a measure of the relative amount of DNA methylation in the amplified region [23]. Comparing PCR amplification at promoter regions upstream of the translation start site of the *INS1*(Fig 7C) and *INS2* (Fig 7D) genes revealed low methylation that was not different between RyR2^KO^, IRBIT^KO^, or control cells. The single CpG site in intron 1 of the *INS1* gene (1-UP4) was extensively methylated but not altered by deletion of RyR2 or IRBIT (Fig 7C). Increased methylation was observed in the proximal portion of exon 2 of *INS1* (Fig 7C). At 1DS3, DNA methylation was increased in IRBIT^KO^ and RyR2^KO^ cells compared to controls. In the 1DS2 region, increased DNA methylation was only observed in the IRBIT^KO^ cells. DNA methylation level in the 1DS2 region is much higher (∼10 fold) compared to the 1-DS1 and 1-DS3 regions. A similar analysis of the *INS2* gene (Fig 7D) showed high methylation at regions downstream of translation start site compared to upstream regions. An increase in DNA methylation was observed in Exon 2 of the *INS2* gene of IRBIT^KO^ and RyR2^KO^ cells compared to controls, with ∼10-fold higher methylation in the 2-DS1 regions compared to the 2-DS2 region. Overall, deletion of RyR2 or IRBIT increased the methylation of CpG sites within exon 2 of both *INS1* and *INS2*, but not in the hypomethylated upstream/promoter regions.

**Fig. 7.**
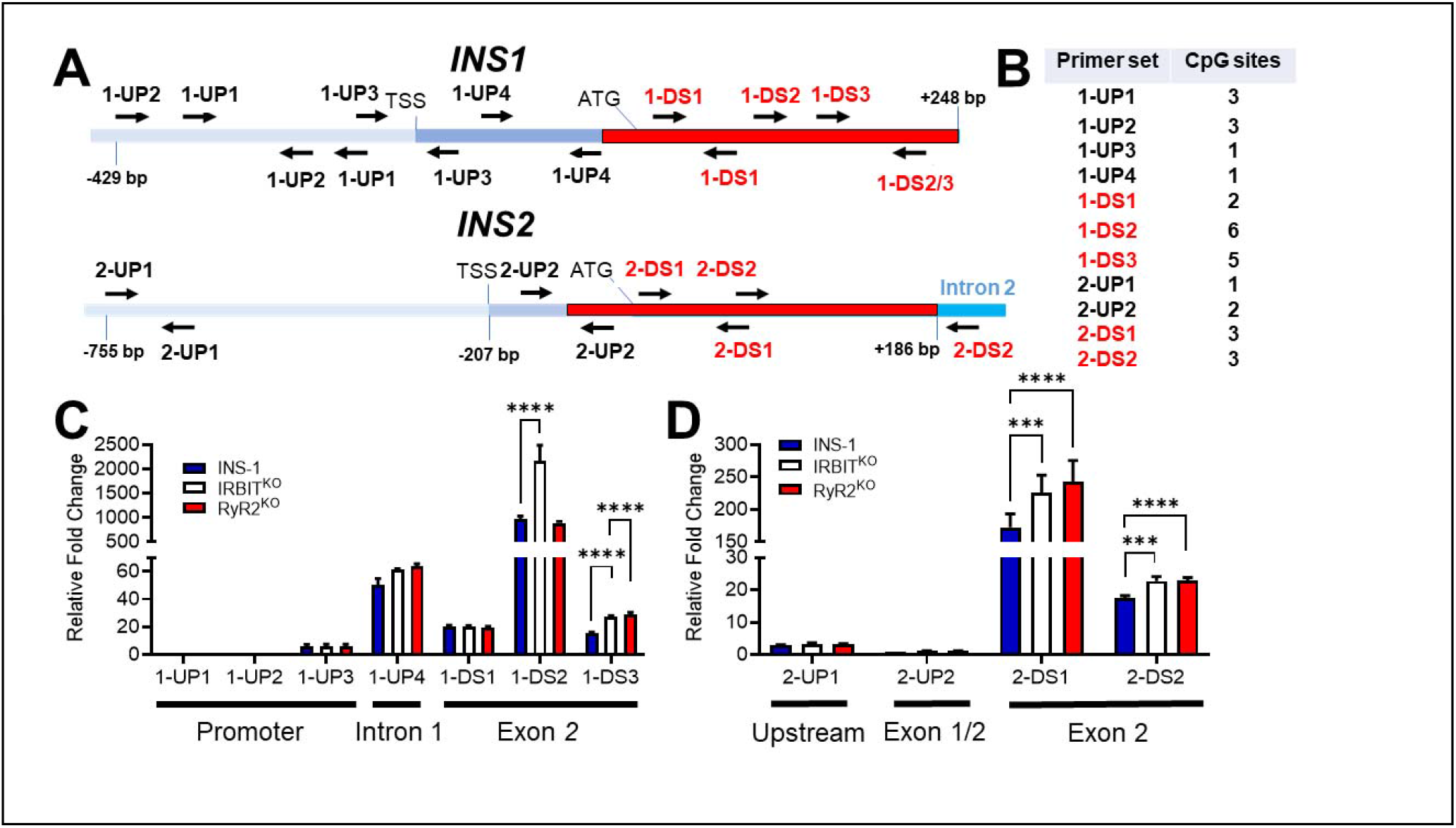
RyR2/IRBIT regulate methylation of the INS1 gene*-*. **A)** Schematic of the *INS1* and *INS2* genes with positions of listed primer pairs. **B)** Number of potential methylation site (CpG) within the sequence amplified by each primer set. **C)** DNA methylation in listed regions of the *INS1* gene in INS-1, IRBIT^KO^, and RyR2^KO^ cells were measured by methylation-dependent qPCR (MD-qPCR). The relative fold changes shown are differences in ΔCq values of the target region normalized to the 1-UP1 region, which stays unmethylated between all cell types. ΔCq is the change in the Cq values for FspEI digested DNA relative to Cq values of the respective undigested DNA. ****, *P* < 0.0001 (two-way ANOVA with Dunnett’s *post-hoc* test). Data shown are the means ± SD for 3 independent experiments. **D)** DNA methylation in indicated regions of the *INS2* gene in INS-1, IRBIT^KO^, RyR2^KO^ and cells was measured by methylation-dependent qPCR (MD-qPCR) as described in **C**. ****, *P* < 0.0001; ***, *P* < 0.001 (two-way ANOVA with Dunnett’s *post-hoc* test). Data shown are the means ± SD for 3 independent experiments.

### Regulation of the INS-1 cell proteome by RyR2 and IRBIT

To assess the effect of RyR2 or IRBIT deletion on the proteome of INS-1 cells, we performed LC-MS/MS analysis of INS-1, RyR2^KO^, and IRBIT^KO^ cells. Deletion of RyR2 resulted in the upregulation of 159 proteins (Fig. S1) and downregulation of 155 proteins (Fig. S2). Deletion of IRBIT resulted in increases levels of 75 proteins (Fig. S3) and decreased levels of 62 proteins (Fig. S4). Of these, 24 proteins were more abundant in both RyR2^KO^ and IRBIT^KO^ cells (Fig 8A) and 17 were less abundant in both RyR2^KO^ and IRBIT^KO^ cells (Fig 8B). GO analysis for overrepresentation of differentially regulated proteins in specific cellular component, biological process, or molecular function categories is shown in Figures 9 and 10. Proteins more abundant in RyR2^KO^ cells were overrepresented in cellular component categories that clustered around synaptic proteins, nuclear proteins, and mitochondrial proteins. Overrepresentation of proteins with increased abundance in biological process (gene expression, cellular nitrogen compound biosynthetic processes, RNA metabolic processes, chromatin organization) and molecular function (RNA binding, mRNA binding, nucleotide catalytic activity, RNA catalytic activity) categories suggest that RyR2 activity may regulate transcription and translation and/or mRNA processing. Proteins more abundance in IRBIT^KO^ cells were overrepresented in three cellular component categories related to mitochondria (mitochondrial matrix, mitochondrial protein-containing complexes, intracellular organelle lumen). Proteins with decreased abundance in RyR2^KO^ cells are mainly overrepresented in categories related to organelle structure and function. Small GTPase binding proteins were overrepresented among proteins with both increased and increased abundance in RyR2^KO^ cells. The smaller group of proteins with decreased abundance in IRBIT^KO^ cells showed modest overrepresentation (< 2-fold) in two cellular component categories, intracellular anatomical structure and cytoplasm.

**Fig. 8.**
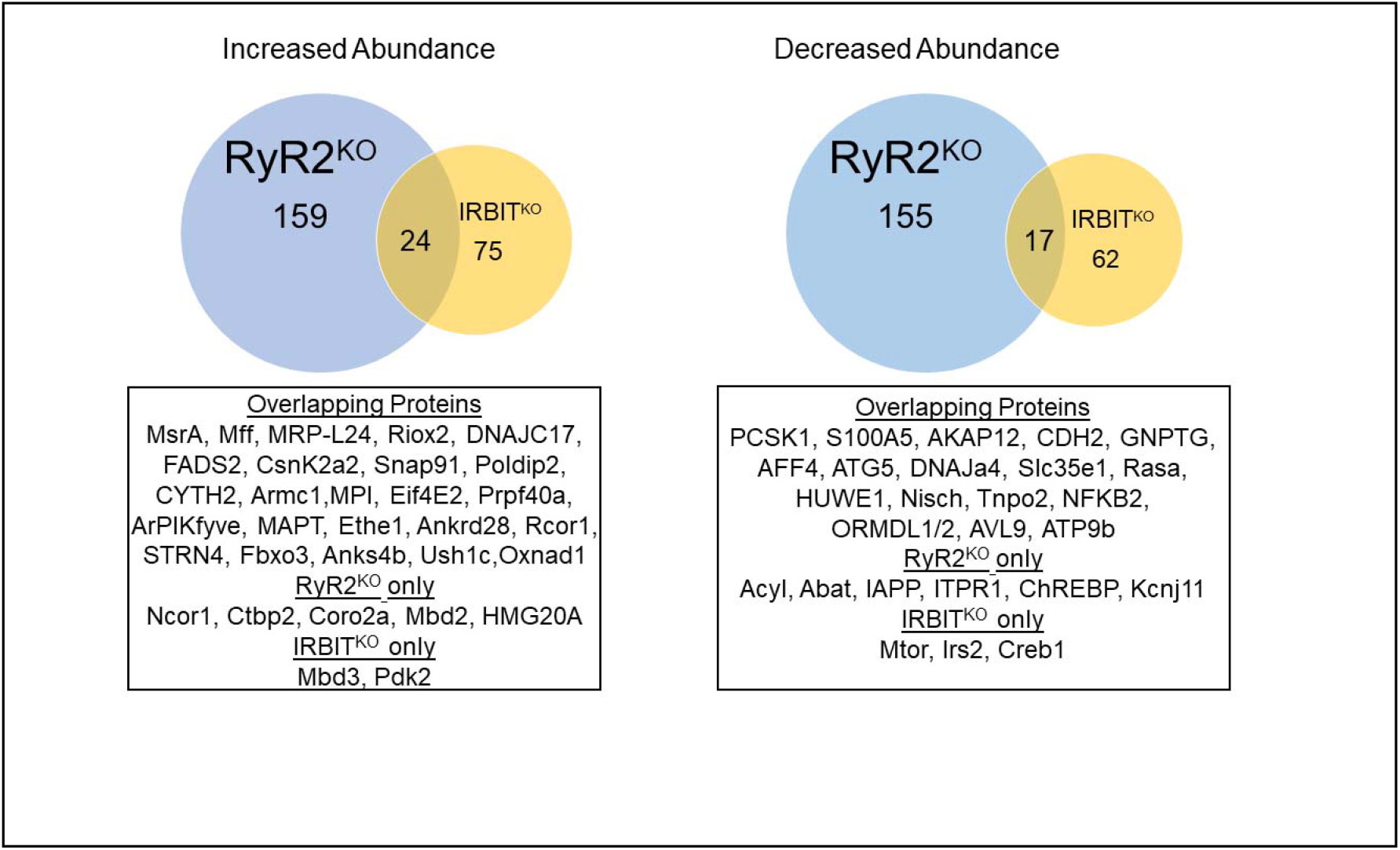
Deletion of RyR2 or IRBIT in INS-1 cells differentially regulates an overlapping set of proteins-. **A)** Venn diagram illustrating the overlap of proteins with increased abundance in RyR2^KO^ and IRBIT^KO^ cells. **B**) Venn diagram illustrating the overlap of proteins with decreased abundance in RyR2^KO^ and IRBIT^KO^ cells. Overlapping proteins are listed in the boxes below each diagram, along with other proteins of interest differentially regulated in either RyR2^KO^ cells or IRBIT^KO^ cells. Protein with decreased abundance were defined as Log_2_(Fold-Change) < -1 and average MS/MS count ratio < 0.5, compared to control. Similarly, proteins with decreased abundance were defined as proteins with Log_2_(Fold-Change) > 1 and average MS/MS count ratio > 2, compared to control. For complete list of differentially regulated proteins see Supplementary Figures 1-4.

**Fig. 9.**
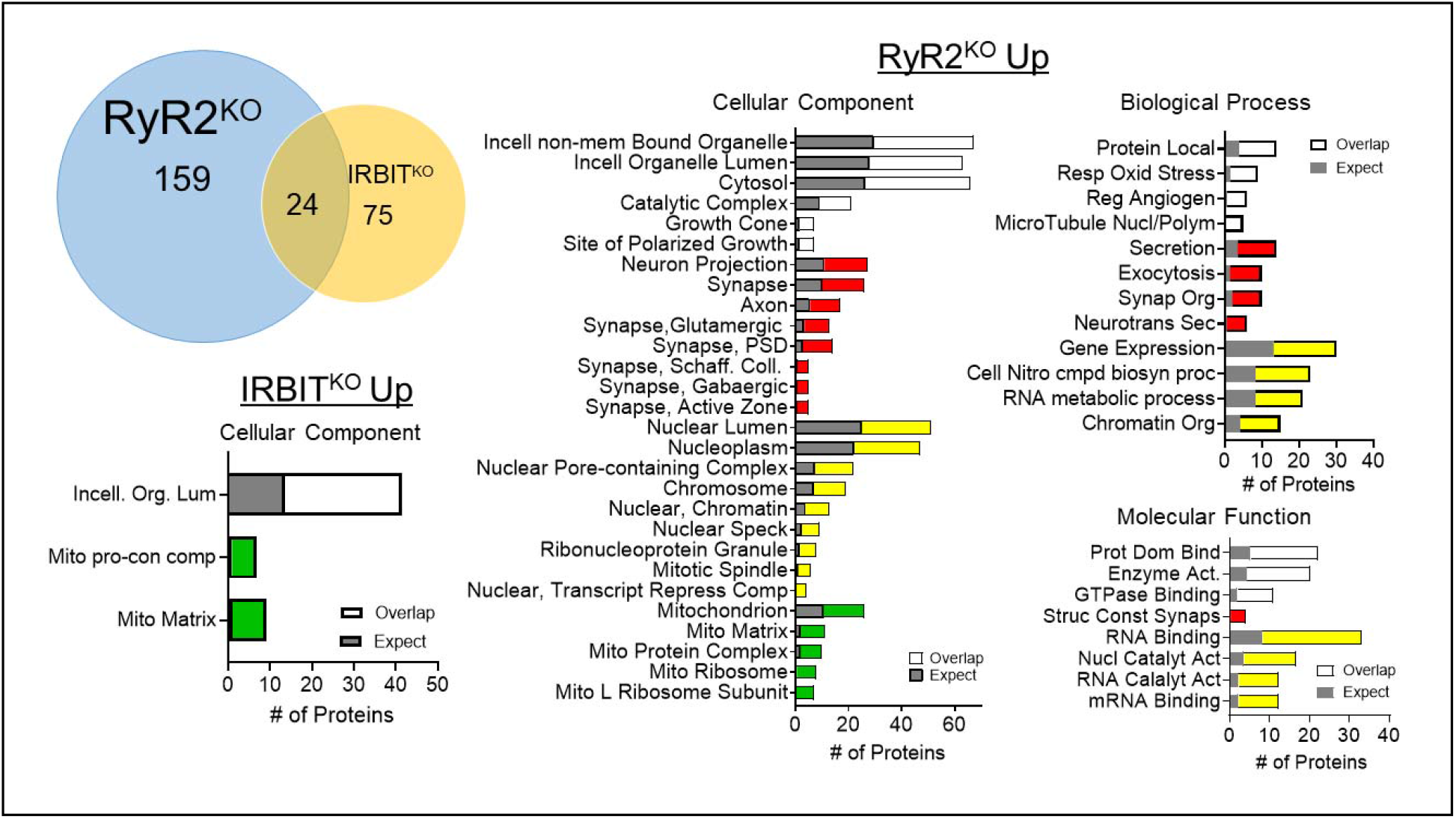
GO analysis of Up-regulated proteins in RyR2^KO^ and IRBIT^KO^ cells-. The number of proteins with increased abundance in RyR2^KO^ or IRBIT^KO^ cells that belong to each of the indicated GO categories (overlap- shown as white, red, yellow, or green bars) is indicated, along with the expected number of proteins in each category (gray bars). Red bars indicate neuronal protein categories, yellow bars indicate nuclear protein categories, and green bars indicate mitochondrial protein categories. FDR < 0.05; overrepresentation > 2-fold.

**Fig. 10.**
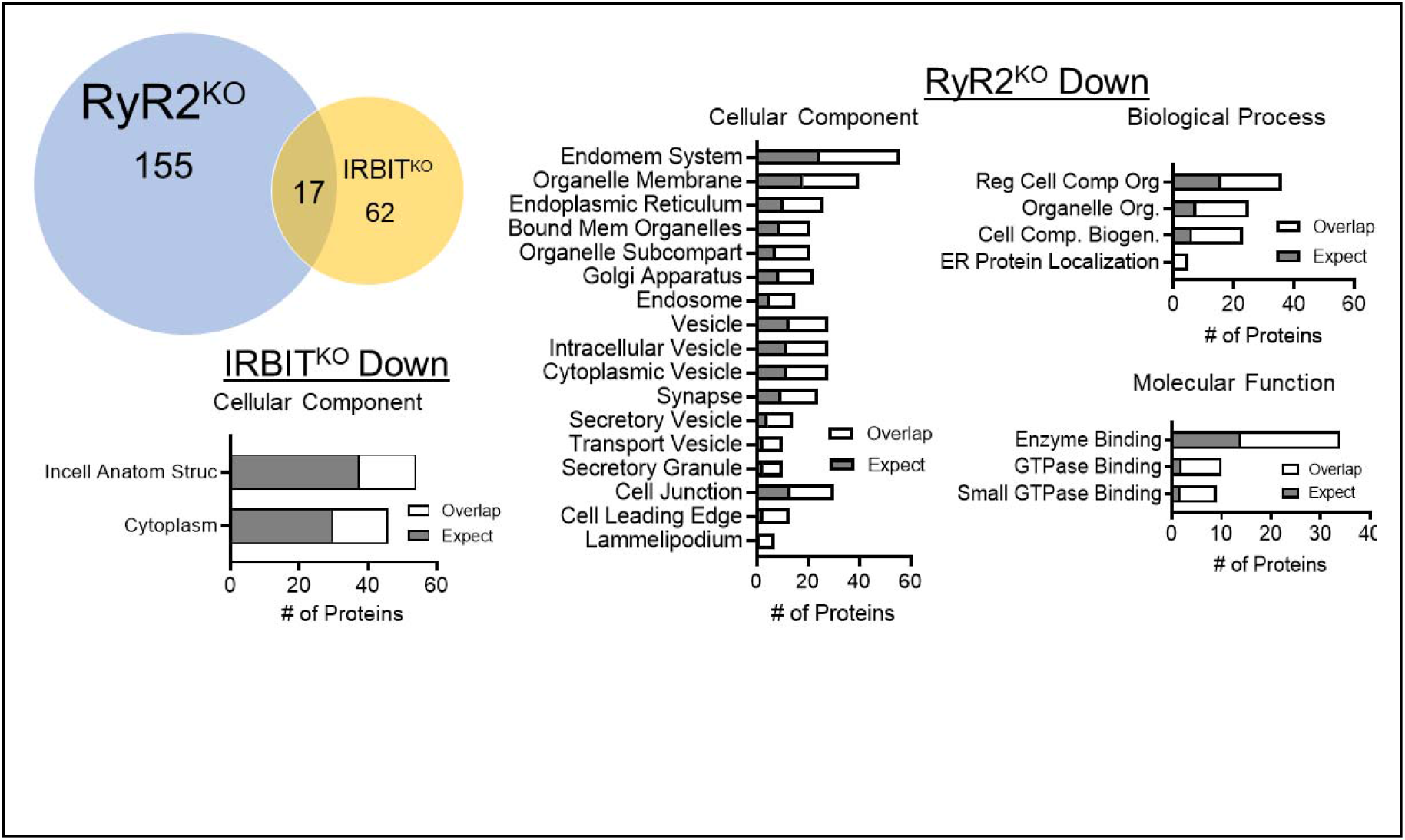
GO analysis of Down-regulated proteins in RyR2^KO^ and IRBIT^KO^ cells-. The number of proteins with decreased abundance in RyR2^KO^ cells that belong to each of the indicated GO categories (overlap- shown as white) is indicated, along with the expected number of proteins in each category (gray bars). FDR < 0.05.

## Discussion

Transcripts for RyR1, RyR2, and RyR3 are detected in INS-1 cells, with RyR2 reported as most abundant [5]. In the current study, deletion of RyR2 essentially abolished mobilization of ER Ca^2+^ by caffeine (Fig 1), suggesting that RyR2 is the predominant, if not sole, functional RyR in INS-1 cells. RyR2 may also contribute to maintenance of resting cytoplasmic [Ca^2+^] (Fig 1F) and PLC activity (Fig 3). Our results suggest a complicated interplay between RyR2 and IP_3_ receptors in INS-1 cells. In control cells, xestospongin C had no effect on the Ca^2+^ integral in response to glucose, but did shorten the period between Ca^2+^ oscillations. In contrast, deletion of RyR2 led to a significant increase in the Ca^2+^ integral in response to glucose, and xestospongin C both decreased the Ca^2+^ AUC and increased the period between oscillations in RyR2^KO^ cells upon glucose stimulation. This result argues that the decrease in IRBIT detected in RyR2^KO^ cells is functionally significant, leading to enhanced IP_3_R activation during glucose stimulation. In both control and RyR2^KO^ cells, the L-type voltage-gated Ca^2+^ channel blocker nicardipine inhibited the Ca^2+^ integral in response to glucose. L-type channels increase intracellular Ca^2+^ levels via conductance of Ca^2+^ into the cell, and via stimulation of RyR2 activity [6]. In the absence of RyR2, L-type channels, via stimulation of PLC activity [24], could facilitate release of Ca^2+^ from disinhibited IP_3_Rs, as well as conductance of Ca^2+^ into the cell via Trp channels [25].

To explain the increased activation of IP_3_ receptors upon RyR2 deletion, we examined PLC activity in response to glucose and the muscarinic receptor agonist carbachol. RyR2 deletion markedly reduced both basal and stimulated IP_1_ accumulation compared to control INS- 1 cells. This apparent decrease in PLC activity wasn’t the result of decreased substrate (i.e. PIP_2_) levels, suggesting that RyR2 plays an important role in supporting PLC activity in INS-1 cells.

Store-operated Ca^2+^ entry (SOCE) plays a key role in supporting PLC activity in pancreatic β-cells [26], and RyR2 is implicated in gating SOCE channels in rat vascular smooth muscle [27]. It will be of interest to determine if a similar mechanism can account for the decreased PLC activity observed in RyR2^KO^ cells.

IRBIT competitively inhibits IP_3_R activation by IP_3_ [14], but also regulates many other proteins [28]. Thus, phenotypes of RyR2^KO^ cells may be attributed to either loss of RyR2 Ca^2+^ release directly, or to downregulation of IRBIT. Deletion of IRBIT had marked effects on Ca^2+^ signaling in the INS-1 cells. Caffeine-stimulated Ca^2+^ transients were maintained in IRBIT^KO^ cells, indicating that RyR2 function is retained in the absence of IRBIT. However, the response was significantly reduced compared to control INS-1 cells. This is likely the result of a reduced pool of Ca^2+^ available for release via RyR2, as the ER Ca^2+^ levels in IRBIT^KO^ cells were reduced compared to control and RyR2^KO^ cells. As expected, deletion of IRBIT led to an increase in the Ca^2+^ response to glucose compared to controls cells, the result of enhanced activation of IP_3_ receptors during glucose stimulation, as in RyR2^KO^ cells. It’s not clear why IRBIT levels/activity are reduced upon RyR2 deletion. We speculate that RyR2 may play a critical role in the phosphorylation of IRBIT on Ser 68 by a Ca^2+^-dependent kinase, leading to further phosphorylation that’s required for IP_3_ receptor binding [15]. Further, unphosphorylated IRBIT is susceptible to proteolytic cleavage [29]. Thus, hypo-phosphorylation of IRBIT could account for both the elevated activity of IP_3_ receptors, and the reduced IRBIT levels observed in RyR2^KO^ cells.

Deletion of either RyR2 or IRBIT had marked effects on insulin secretion, cellular insulin content, and insulin gene transcript levels. RyR2 deletion had the greatest effect on basal (2.5 mM) and glucose stimulated insulin secretion, but IRBIT deletion also reduced basal and glucose-stimulated insulin secretion compared to control INS-1 cells. Although elevated Ca^2+^ release from IP_3_ receptors during glucose stimulation was a hallmark of both RyR2 and IRBIT deletion, xestospongin C didn’t inhibit insulin secretion in either control, RyR2^KO^, or IRBIT^KO^ cells. Our results show that even when IP_3_ receptors are highly active, they don’t contribute to glucose-stimulated insulin secretion in INS-1 cells. This is consistent with previous reports that IP_3_ receptors are upregulated in the β-cells of type 2 diabetics, but associated with impairment of glucose-stimulated insulin secretion and β-cell dysfunction [12].

To understand what might account for this reduction in insulin secretion, we examined the insulin content of control INS-1, RyR2^KO^, and IRBIT^KO^ cells and found an ∼70% decrease in insulin content of RyR2^KO^ cells, and an ∼40% decrease in insulin content in IRBIT^KO^ cells compared to controls. This decrease in insulin content was accompanied by a similar decrease in *INS2* gene transcript levels in both RyR2^KO^ and IRBIT^KO^ cells. Interestingly, the level of *INS1* transcript detected in RyR2^KO^ cells was reduced compared to controls, but wasn’t different in IRBIT^KO^ cells. Although the role of IRBIT as a regulator of IP_3_ receptor activation is well studied, it’s clear that IRBIT plays many other roles [30]. IRBIT contains a highly conserved, but catalytically inactive, S-adenosyl homocysteine hydrolase (AHCY) domain [31]. By virtue of this domain, IRBIT can bind to and may regulate the activity of catalytically active AHCY [21] as well as its distribution between the cytoplasm and the nucleus [22]. Since AHCY degrades S-adenosyl homocysteine (SAH), a potent inhibitor of DNA methyltransferases [32], dysregulation of AHCY by loss of IRBIT could play a role in the increased *INS1* and *INS2* gene methylation that we observed in RyR2^KO^ and IRBIT^KO^ cells. While deletion of RyR2 slightly increased nuclear AHCY localization, deletion of IRBIT caused a marked shift of AHCY from the cytoplasm into the nucleus, though the total amount of AHCY immunostaining detected in the nuclei of IRBIT^KO^ cells was reduced compared to controls. Increased AHCY activity in the nucleus could disinhibit DNA, RNA, and protein methyltransferases (Fig 6D). AHCY binds to chromatin near transcription start sites of active genes [33], suggesting that AHCY regulation of DNA methylation may be spatially specific. Given that AHCY is only active as a homotetramer [34], incorporation of IRBIT via its AHCY domain into AHCY complexes could diminish AHCY activity. We speculate that, in addition to controlling AHCY nuclear accumulation, IRBIT may also limit AHCY activity. Such a scenario would imply that AHCY in the absence of IRBIT not only accumulates preferentially in the nucleus, but may be catalytically more active.

The shift of AHCY from the cytoplasm to the nucleus in IRBIT^KO^ cells and the corresponding changes in insulin mRNA levels were coincident with an increase in methylation of both the *INS1* and *INS2* genes downstream of the translation start site. The promoter/upstream regions were hypomethylated in both *INS1* and *INS2*, which didn’t change upon deletion of either RyR2 or IRBIT, consistent with the finding that hypomethylation of the promoter regions of insulin genes serves as a marker of islet cell identity [35]. Methylation of genes within the first exon (or within ∼200 bp downstream of the transcription start site) is highly correlated with inhibition of transcription [36]. The increased methylation in *INS1* and *INS2* genes in RyR2^KO^ and IRBIT^KO^ cells was observed in exon 2. Studies in pancreatic islets from NOD mice found an inflammation-mediated increase in *INS1* exon 2 and *INS2* exon 1 methylation and a corresponding decrease in *INS1 and INS2* mRNA [37]. The same study found no correlation between methylation of the *INS2* promoter region and *INS2* mRNA levels. The human insulin gene is hypermethylated at the TSS +63 position, corresponding to the rat *INS2* 2-UP2 region, in type 2 diabetic patients, and this hypermethylation is correlated with decreased *INS* mRNA [38]. However, we observed extremely low methylation in this region in both RyR2^KO^, IRBIT^KO^, and control cells. While it’s possible increased methylation in exon 2 of *INS2* in RyR2^KO^ and IRBIT^KO^ cells contributes to the decreased *INS2* mRNA levels, that conclusion is not currently supported.

The differential effect of RyR2 or IRBIT deletion on *INS1* mRNA levels was also coincident with differential changes in methylation of the *INS1* gene in exon 2. While CpG site(s) in exon 2 (1-DS3) are hypermethylated in both RyR2^KO^ and IRBIT^KO^ cells, a specific CpG site in exon 2 (ATG +136) in the *INS1* gene is hypermethylated in IRBIT^KO^ cells compared to control and RyR2^KO^ cells. It is currently unclear whether increased methylation within the 1- DS3 region accounts for the decrease in INS1 mRNA in RyR2^KO^ cells, or if the specific increase in methylation at +136 is responsible for the maintenance of *INS1* mRNA levels in IRBIT^KO^ cells. However, this differential regulation of *INS1* mRNA levels may account for the smaller decrease in insulin content in IRBIT^KO^ cells, compared to RyR2^KO^ cells. It will be of interest to determine if RyR2 or IRBIT deletion results in a more global increase in DNA methylation.

Deletion of RyR2 or IRBIT differentially regulates a partially overlapping set of proteins. It’s not clear if this regulation is occurring pre- or post-translationally, but the reduction of IRBIT protein in RyR2^KO^ cells, with no decrease in mRNA levels, demonstrates that RyR2 activity is capable of regulating protein levels post-transcriptionally. GO analysis revealed that RNA binding/processing proteins are overrepresented in the population of proteins increased in abundance by RyR2 deletion, suggesting altered RNA processing in the absence of RyR2.

Mitochondrial proteins are also overrepresented in the population of proteins increased in abundance by RyR2 or IRBIT deletion, perhaps reflecting the proposed role of IRBIT in regulating Ca^2+^ flux between the ER and mitochondria [39]. The decrease in ATG5 protein in both RyR2^KO^ and IRBIT^KO^ cells might also explain the increased levels of some mitochondrial proteins, since deletion of ATG5 (autophagy-related gene 5) in T-lymphocytes was shown to differentially regulate mitochondrial protein levels and mitochondrial mass [40]. GTPase binding proteins are overrepresented in populations of both increased and decreased abundance proteins in RyR2^KO^ cells, suggesting a switch in the complement of modulators of small GTPase proteins, which play critical roles in vesicle trafficking [41]. Finally, proteins more abundant upon RyR2 deletion are overrepresented in several categories related to the nucleus, including four components of transcription repressor complexes- Rcor1 (REST co-repressor 1), Ncor1 (nuclear receptor corepressor 1), Ctbp2 (c-terminal binding protein 2), and Coro2a (coronin 2a). Rcor1, also increased in abundance in IRBIT^KO^ cells, and Ctbp2 are of particular interest since they are part of the RE-1 Silencing Transcription factor (REST) repressor complex [42], which represses genes critical for β-cell function, but is inactivated during differentiation. Several proteins repressed by REST (Pcsk1, neuroendocrine convertase 1; Chga, chromogranin A; Chgb, secretogranin; Stmn2, stathmin 2) [42] are reduced in abundance in RyR2^KO^ cells. It may be of interest to examine the possibility that loss of RyR2 function permits expression of some components of the REST repressor complex, and thus repression of a subset of genes critical for β-cell function.

Some of the differentially regulated proteins identified in RyR2^KO^ and IRBIT^KO^ cells are dysregulated in diabetes and/or play a key role in β-cell function. Among these are PCSK1 [43] and ORMDL1/2 (sphingolipid biosynthesis regulator 1/2) [44] which are reduced in both RyR2^KO^ and IRBIT^KO^ cells, and IAPP (islet amyloid polypeptide), Acyl (ATP-citrate lyase), and Abat (GABA aminotransferase) [45], and Kcnj11 (Kir6.2) [46] which are reduced in RyR2^KO^ cells. The Ca^2+^-dependent adhesion molecule Cdh2 (N-cadhedrin) is also reduced in both RyR2^KO^ and IRBIT^KO^ cells. Cell adhesion and spreading via N-cadhedrin enhances GSIS [47].

The pancreas-specific deletion of HUWE1 (HECT, UBA, and WWE domain containing E3 ubiquitin ligase 1), which is reduced in both RyR2^KO^ and IRBIT^KO^ cell, leads to increased β-cell apoptosis and reduced β-cell mass [48]. Anks4b which is, together with its binding partner Ush1c (Harmonin), more abundant in both RyR2^KO^ and IRBIT^KO^ cells, increases susceptibility to ER stress-induced apoptosis when overexpressed in MIN6 cells [49]. Activation of the imidazole receptor NISCH (nischarin), which is reduced in both RyR2^KO^ and IRBIT^KO^ cells, potentiated GSIS and stimulated proliferation of INS-1 cells [50]. Finally, some key proteins in β-cell function were specifically reduced in IRBIT^KO^ cells, including mTOR (mammalian target of rapamycin) [51], Creb1 (cAMP response element-binding protein 1) [52], and IRS2 (insulin receptor substrate 2) [53].

In summary, deletion of RyR2 in INS-1 cells had the unanticipated effect of reducing IRBIT levels and activity. Deletion of RyR2 or IRBIT both enhanced IP_3_ receptor activation during glucose stimulation, and reduced insulin secretion, content, and *INS2* mRNA. In addition, the *INS1* and *INS2* genes were hypermethylated in exon 2 upon RyR2 or IRBIT deletion, coincident with alterations in the nuclear localization of AHCY. One limitation of this study is that, while insulin content is clearly reduced by RyR2 or IRBIT deletion, it’s not clear if this reduction accounts for the reduced basal or glucose-stimulated insulin secretion, since granule trafficking/exocytosis may also be impaired by RyR2 or IRBIT deletion. Furthermore, the relationship between RyR2 activity and IRBIT levels remains unknown. Nevertheless, IRBIT regulation of AHCY localization and, potentially, activity positions it to regulate the activity of protein, RNA, and DNA methyltransferases via modulation of local SAH levels, and regulate the proteome. It will be of interest to determine if pathophysiological perturbation of RyR2 activity or dysregulation of ER Ca^2+^ levels in pancreatic β-cells leads to dysregulation of IRBIT activity.

## Materials and Methods

### Chemicals and Reagents

Antibodies to pan-specific RyR, AHCY, AHCYL1, and β-actin, and mouse IgG-κ binding protein conjugated to CFL 488 were from Santa Cruz Biotechnology (Dallas, TX). Goat anti-mouse IgG conjugated to IRDye 680RD and goat anti-rabbit IRDye 800CW were from LI-COR (Lincoln, NE). Antibodies to phosphatidylinositol 4,5 bisphosphate (PIP_2_) were from Echelon Biosciences (Salt Lake City, UT). Goat anti-mouse IgG conjugated to horseradish peroxidase was from BioRad (Hercules, CA). ERK1/2 and pERK1/2 antibodies were from Cell Signaling Technology (Danvers, MA).

Oligonucleotides encoding gRNA and primers were obtained from Integrated DNA Technologies (Coralville, IA). The pSpCas9(BB)-2A-Puro (PX459) V2.0 vector was a gift from Feng Zhang (Addgene plasmid # 62988). T4 polynucleotide kinase and T7 DNA ligase were from New England Biolabs (Ipswich, MA). Fast AP thermosensitive alkaline phosphatase and FastDigest Bpil were from Thermo Scientific (Waltham, MA). Plasmid-Safe ATP-Dependent DNase was from Lucigen (Middleton, WI). Surveyor Mutation Detection Kit S100 was from Integrated DNA Technologies (Coralville, IA).

*Chemicals-* Fura-2 AM was from Invitrogen (Carlsbad, CA). Xestospongin C was from Cayman Chemical (Ann Arbor, MI). All other reagents, unless otherwise indicated, were from Sigma-Aldrich (St. Louis, MO).

### Cell Culture

INS-1 cells (Gift of Dr. Ming Li, Tulane University) were cultured in RPMI-1640 medium (Sigma-Aldrich) supplemented with 10% fetal bovine serum (Qualified,Gibco), 11 mg/mL sodium pyruvate, 10 mM HEPES, 100 U/mL penicillin, 100 μg/mL streptomycin, and 50 μM mercaptoethanol at 37°C, 5% CO_2_.

### Construction of Cas9 Plasmids

gRNA sequences were designed using the crispr.mit.org website. Oligonucleotides were synthesized by IDT (Coralville, IA).

RyR2 gRNA (Exon 6): 89-Forward 5’-CACCGTTTGTCGGTGGAAGACCGGG-3’

89-Reverse 5’-AAACCCCGGTCTTCCACCGACAAA-3’

92-Forward 5’-CACCGCCGGTCTTCCACCGACAAAC-3’

92-Reverse 5’-AAACGTTTGTCGGTGGAAGACCGG-3’

RyR2 Exon 6 primers: Forward 5’-GTGGAAATCAGTGCGGAGTC-3’

Reverse 5’-TGTATTTGGGTTCTGCAAAGG-3’

AHCYL1 gRNA (Exon 6): 75-Forward 5’-CACCGCATTGACCGCTGTGTCAACA-3’

75-Reverse 5’-AAACTGTTGACACAGCGGTCAATG-3’

AHCYL1 Exon 6 primers: Forward 5’-GAGGCATCTGTTGCTGTTCA-3’

Reverse 5’-CTCCAGCATTCCTGCTTCAG-3’

gRNA oligonucleotides were subcloned into pSpCas9(BB) using BbsI (New England Biolabs). Ligation products were used to transform competent DH5α *E. coli,* and transformants were selected on Luria broth-agar plates containing 100 µg/mL ampicillin. Plasmid DNA was purified and sequenced (Purdue Genomics Core Facility) to confirm assembly of the desired construct.

### Generation and Validation of Knockout Clones

INS-1 cells were transfected with 2 μg RyR2- or IRBIT-targeted gRNA/Cas9 plasmid or pEGFP-N1 (selection control) using Lipofectamine 2000 (Invitrogen) per manufacturer’s instructions. 72 hours post-transfection, cells were selected with 3 μg/mL puromycin until no cells remained in the pEGFP-N1 transfected well. Individual clones from the gRNA/Cas9 transfected cells were then isolated by limiting dilution in 96-well plates (Corning). RPMI-1640 media was changed weekly for 4 weeks, and clones were gradually expanded. Once expanded, clones were plated at 90% confluency in 96-well plates and allowed to incubate overnight at 37°C, 5% CO_2_. Cells were lysed and genomic DNA was extracted using QuickExtract DNA Extraction Buffer (Lucigen) per the manufacturer’s instructions. Extracted genomic DNA from INS-1, RyR2^KO^, or IRBIT^KO^ cells was subjected to PCR amplification (Herculase II Fusion DNA Polymerase and 5X Herculase II PCR Buffer and dNTP (Agilent Techologies) using primers flanking the region targeted by the gRNA. Purified amplicons were sequenced at the Purdue University Genomics Core.

### Single-Cell Intracellular Ca^2+^ Assays

INS-1 and RyR2^KO^ cells were either plated in a 35 mm tissue culture dish (Corning) containing a poly-D-lysine coated round glass coverslip (for assays using perfusion; Warner Instrument) or plated in a poly-D-lysine coated 4-chambered 35 mm glass bottom tissue culture dish (for assays not using perfusion; Cellvis). Cells were incubated overnight in RPMI-1640 media at 37°C, 5% CO_2_. For glucose assays, cells were deprived of glucose for an additional 24 hours in low glucose RPMI-1640 media. Cells were washed twice with PBS prior to loading with 3 μM of the Ca^2+^ indicator Fura-2 AM (Invitrogen) diluted in a modified Krebs-Ringer buffer [KRBH: 134 mM NaCl, 3.5 mM KCl, 1.2 mM KH_2_PO_4_, 0.5 mM MgSO_4_, 1.5 mM CaCl_2_, 5 mM NaHCO_3_, 10 mM HEPES (pH 7.4)] supplemented with 0.05% fatty acid free BSA at room temperature for 1 hour. The KRBH containing Fura-2 AM was then removed, and the cells were washed twice with KRBH, then equilibrated for 30 minutes at room temperature in KRBH alone or KRBH containing a 2x concentration of indicated inhibitors. For perfusion assays, the glass coverslip was mounted on a perfusion chamber attached to the stage of an Olympus IX50 inverted microscope equipped with a PlanApo 40x objective lens (0.95 na) and solutions/stimuli were perfused to the chamber at a constant flow rate (1 mL/min) at room temperature. For assays not using perfusion, the 4-chambered 35 mm dish was mounted on a chamber attached to the stage of the microscope. Cells were stimulated with the indicated stimulus at a 2x concentration. Cells were alternatively excited at 340/11 nm and 380/20 nm wavelengths using a band pass filter shutter (Sutter Instrument) and changes in intracellular Ca^2+^ were measured by recording the ratio of fluorescence intensities at 508/20 nm in time lapse (time interval of 0.6 seconds) using a Clara CCD camera (Andor Technology). Background subtraction from the raw 340/11 nm and 380/20 nm wavelengths was performed, then isolated single cells were selected as regions of interest (ROI) and the 340/11 nm/380/20 nm ratios for each ROI were measured using MetaMorph image analysis software (Molecular Devices). All single-cell Ca^2+^ transients were normalized to their baseline intracellular Ca^2+^ level, which was obtained by averaging the 340/11 nm/380/20 nm ratios during the first minute of each experiment when no stimulus was present. Ca^2+^ transients are plotted as normalized 340/11 nm/380/20 nm ratios against time.

### Intracellular Ca^2+^ Measurements in 96-well Plates

INS-1 and RyR2^KO^ cells were plated at 70-90% confluency in black-walled 96-well plates (Corning) in RPMI-1640 media and incubated overnight at 37°C, 5% CO_2_. Cells were washed twice with PBS and incubated with 100 μL 5 μM Fura-2 AM in KRBH for 1 hour at room temperature. The KRBH containing Fura-2 AM was removed, the cells were washed twice with KRBH, and equilibrated for 30 minutes in 100 μL KRBH at room temperature. Cells were stimulated by injection of 100 μL 10 mM caffeine (2x) or KRBH (buffer control). Changes in intracellular Ca^2+^ concentrations were measured by recording the ratio of fluorescence intensities at 508/20 nm resulting from excitation of Fura-2 AM at 340/11 nm or 380/20 nm (center/bandpass) using a Synergy 4 multimode microplate reader (BioTek). Ratios were acquired every 0.7 seconds for 15 seconds before injection and 2 minutes after injection. Data were corrected for injection artifact by subtracting the change in fluorescence ratio measured in cells injected with KRBH alone.

### IP_1_ HTRF Assays

INS-1 and RyR2^KO^ cells were plated at approximately 200,000 cells/well in an opaque 96-well tissue culture plate (Corning) and incubated overnight in low glucose RPMI- 1640 media at 37°C, 5% CO_2_. Cells were washed with PBS and incubated in a pre-stimulation buffer [10 mM HEPES, 1 mM CaCl_2_, 0.5 mM MgCl_2_, 4.2 mM KCl, 146 mM NaCl (pH 7.4)] for 1 hour at 37°C, 5% CO_2_. The pre-stimulation buffer was decanted, and stimulants and/or inhibitors at the concentrations indicated, were applied in the same buffer supplemented with 50 mM LiCl to inhibit inositol monophosphate degradation were added to the cells and incubated for 1 hour at 37°C, 5% CO_2_. Accumulation of IP_1_ was measured using the IP-One Gq Homogenous Time-Resolved Fluorescence (HTRF) kit from Cisbio per the manufacturer’s instructions. The IP_1_ concentration of each sample was interpolated by comparison to a standard curve of IP_1_ concentrations.

### Endoplasmic Reticulum Ca^2+^ Measurements

INS-1 and RyR2^KO^ cells were plated in a 6-well dish (Corning) in RPMI-1640 media at 70-90% confluency and transfected with 1 μg DNA encoding the endoplasmic reticulum Ca^2+^ indicator pcDNA-D1ER [19] using Lipofectamine 2000 per the manufacturer’s instructions. 24 hours after transfection, cells were transferred to a 4-chambered 35 mm glass bottom dish (Cellvis) coated with poly-D-lysine. 16-24 hours prior to imaging, cells were incubated in low glucose RPMI-1640 media overnight at 37°C, 5% CO_2_.D1ER FRET measurements were performed on a Nikon A1 confocal microscope using a 20x objective. The CFP-YFP FRET pair was excited with a 457 nm argon laser line, and CFP and YFP (FRET) emissions were collected using 482/35 nm and 525/25 nm PMT filters, respectively, before and after treatment of cells with 1 µM thapsigargin for 30 minutes.

### Insulin Assays

INS-1 and RyR2^KO^ cells were plated at 70-90% confluency in 24-well plates (Corning) in RPMI-1640 media and incubated overnight at 37°C, 5% CO_2_. 16-24 hours prior to assay, cells were incubated in serum-free, low glucose RPMI-1640 media supplemented with 0.1% fatty acid-free BSA overnight at 37°C, 5% CO_2_. Cells were washed once with PBS and pre-incubated with 1 mL fatty acid-free KRBH alone or containing the working concentration of inhibitors for 30 minutes at 37°C, 5% CO_2_. After 30 minutes, KRBH was removed and replaced with either 1 mL KRBH or KRBH containing the indicated concentrations of stimulants, and cells were stimulated for 30 minutes at 37°C, 5% CO_2_. Supernatants were collected and stored at -20°C until assayed. Cells were lysed in 200 μL ice-cold RIPA lysis buffer supplemented with protease inhibitors (1 mM 4-(2-aminoethyl) benzenesulfonyl fluoride hydrochloride, 800 nM aprotinin, 50 μM bestatin, 15 μM E-64, 20 μM leupeptin, and 10 μM pepstatin A) and lysates were transferred to 1.5 mL tubes on ice. After 20 minutes, lysates were clarified by centrifugation (14,000 rcf for 10 minutes at 4°C) and supernatants were transferred to new tubes. Protein content was measured using the Pierce BCA Protein Assay Kit (Thermo Fisher) per the manufacturer’s instructions. Insulin measurements were performed either using High-Range insulin ELISA (Alpco) or Insulin High-Range assay kits (Cisbio). For insulin content assays, cells were extracted with 70% ethanol/0.4M HCL for 12 hours at 4°C, then neutralized with 1 M Tris pH 8.0, (20-fold final dilution) before insulin and protein were assayed as described above.

### Immunoblotting

INS-1, RYR2^KO^, or IRBIT^KO^ cells were plated at 70-90% confluency in 6-well plates (Corning) in RPMI-1640 media and incubated overnight at 37°C, 5% CO_2_. The following day, cells were washed once with ice-cold PBS and lysed in 200 μL TBS containing 1% Triton- X100 supplemented with protease inhibitors. Cells lysates were incubated on ice for 30 minutes and then transferred to 1.5 mL tubes to be clarified by centrifugation (14,000 rcf for 10 minutes at 4°C). Protein concentrations were determined using the BCA Protein Assay kit (Thermo Fisher). 30 μg of each lysate was separated by SDS-PAGE on 8% acrylamide gels at 150 V for 1 hour. Proteins were transferred onto PVDF membranes in ice-cold Towbin Buffer (25 mM Tris (pH 8.3),192 mM glycine,10% ethanol) at 100V for 1 hour. Membranes were blocked in 5% non-fat milk in TBS with 0.1% Tween 20 (TBST) for 1 hour at room temperature. Blocked membranes were incubated with primary antibody (1:1000) in 5% milk in TBST overnight at 4°C. The following day, membranes were washed 3 times with TBST and were then incubated with secondary antibodies (IRDye 680RD goat anti-mouse IgG (1:10,000) or goat anti-mouse IgG HRP (1:10,000)) in 5% milk in TBST for 1 hour at room temperature. Blots were imaged on a LI-COR Odyssey CLx imager and analyzed using Image Studio (IRDye 680RD) or an Azure Biosystems Sapphire imager (chemiluminescence). For RYR blots, crude microsomes were isolated using the Endoplasmic Reticulum Isolation Kit (Sigma) per the manufacturer’s instructions. 4-6 confluent 15 cm dishes worth of cells were harvested by trypsin for each cell line prior to microsome collection. All buffers were supplemented with protease inhibitors immediately prior to use. Crude microsomal pellets were resuspended in isotonic buffer (10 mM HEPES, pH 7.8, with 0.25 M sucrose,1 mM EGTA, and 25 mM potassium chloride) supplemented with protease inhibitors and homogenized by repeatedly passing through a 23g needle. Protein concentrations were determined by Bradford Assay (Thermo). Prior to electrophoresis, microsomes were heated to 65°C for 10 min in Laemmeli sample buffer supplemented with 5% mercaptoethanol. 90-180 ug either INS-1 or RYR2^KO^ microsomes were separated by SDS-PAGE on 5% acrylamide gels at 75 V for 3 hours. Proteins were transferred onto PVDF membranes in ice-cold Towbin Buffer, 10% methanol at 25V for 16 hour at 4°C . Membranes were blocked in 3% BSA in TBST for 1 hour at room temperature and then incubated with anti-RyR (F-1) (1:1000) in 3% BSA in TBST overnight at 4°C. Primary antibody was removed the following day, and membranes were incubated in goat anti-mouse IgG HRP (1:10,000)) in 5% milk in TBST for 1 hour at room temperature. Chemiluminescence was detected using standard ECL reagents and imaged on Azure Biosystems Sapphire imager.

### RT-qPCR Assays

Total RNA was extracted from the indicated cell lines using TRIzol (Thermo Fisher). 1 μg of total RNA was used to transcribe cDNA using a high-capacity RNA-to-cDNA kit (Applied Biosystems) per the manufacturer’s protocol. 10 ng of each cDNA was used to perform RT-qPCR using the PowerTrack SYBR Green Master Mix (Thermo Fisher) in a Viia7 real-time PCR system (Applied Biosystems). The protocol for RT-qPCR was as follows: initial activation at 95°C for 5 minutes was followed by 45 cycles of denaturation at 95°C for 10 seconds, primer annealing at 62°C for 30 seconds, and extension at 72°C for 30 seconds, followed by a melting curve step. Cycle threshold (Ct) values were determined for each primer pair, and ΔCT values were separately determined for target mRNAs against two different reference genes, glyceraldehyde 3-phosphate dehydrogenase (GAPDH) and phosphoglycerate kinase 1 (PGK1). ΔΔCT values were calculated for each target mRNA in RyR2^KO^ and IRBIT^KO^ cell lines relative to control INS-1 cells and were used to determine fold difference (2^-ΔΔCT^). The oligonucleotides used in rt-qPCR experiments were designed using Primer-Blast online tool (https://www.ncbi.nlm.nih.gov/tools/primer-blast/).

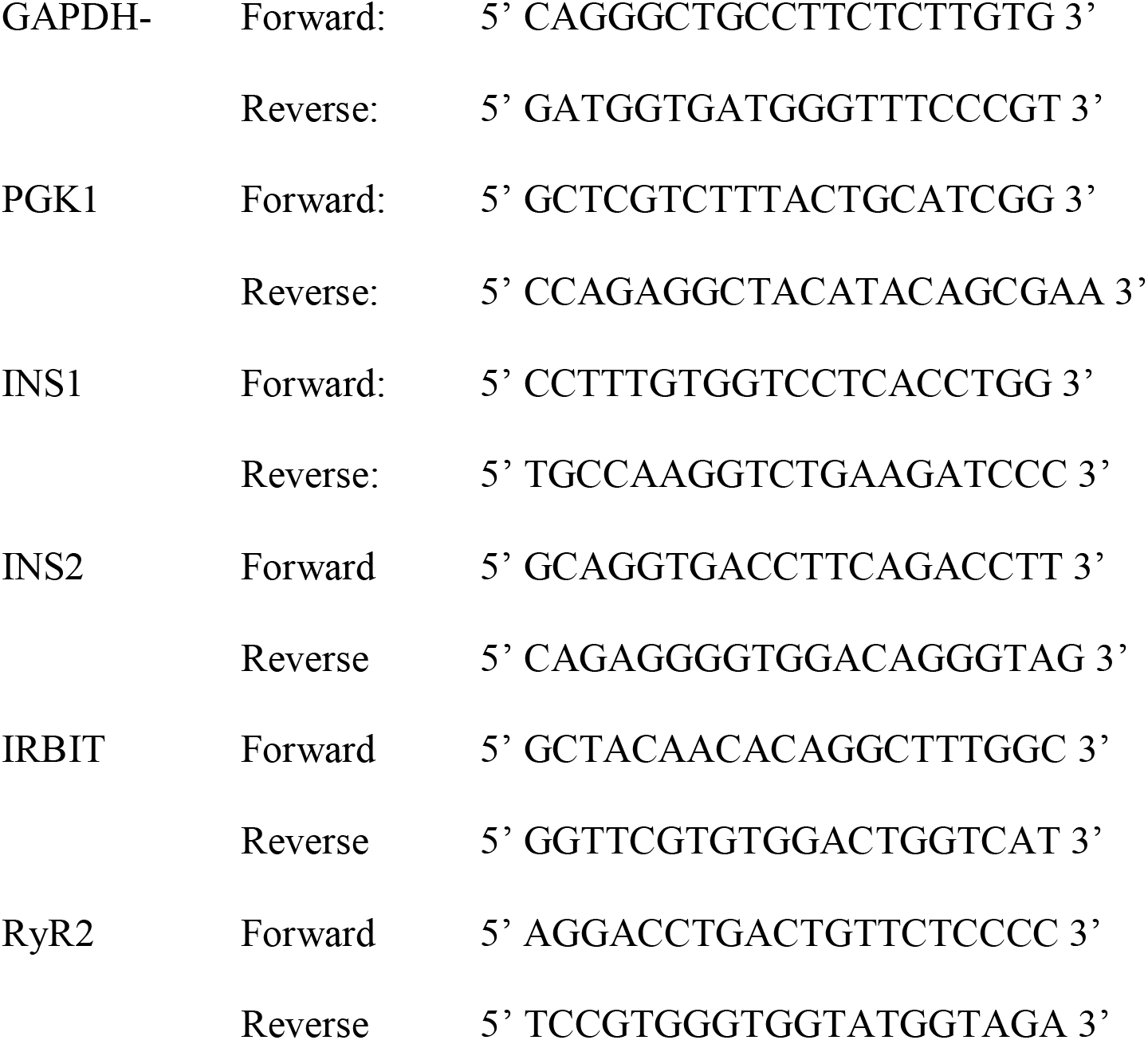

### DNA Methylation-Dependent qPCR Assay (MD-qPCR)

Genomic DNA was isolated from cells using standard phenol: chloroform isolation protocol, followed by ethanol precipitation. The DNA was RNase treated and purified again. 14 μg of sample DNA were digested overnight at 25°C with CviQI restriction enzyme (NEB, R0639L), which cuts outside the region of interest.

The next day, the samples were purified, and 5 μg were digested overnight at 37°C with FspEI (NEB, R0662S), which recognizes C^M^C sites and creates a double-stranded DNA break on the 3’ side of the modified cytosine at N12/N16. The purified CviQI and CviQI+FspEI digested DNA were quantified by PicoGreen according to the manufacturer’s protocol (Life Technologies, P11495) using a NanoDrop 3300 fluorescence spectrophotometer. The digested DNA was checked on an agarose gel to check the digestion of the samples. The quantitative PCR was performed using 6 ng of singly cut (CviQI only) and doubly cut (CviQI+FspEI) DNA for each sample, using the qPCR master mix EvaGreen according to the manufacturer’s conditions (MidSci, BEQPCR-S). The change in DNA methylation is represented by the relative fold change in the Cq value as follows: 2^(ΔCq(S)- ΔCq(C)), where ΔCq is the Cq change in (CviQ1+FspEI) – (CviQ1) digested sample. C is the normalization control region, 1-UP1, which does not methylate, and S is the target region to check the methylation. Standard deviations represent three technical and two biological replicates. An increase in Cq value indicates a gain of DNA methylation. The primers were designed for *INS1* and *INS2* promoter region and their integrity was validated by assessing the size of PCR products on a polyacrylamide gel. The oligonucleotides used in the MD-qPCR experiments were:

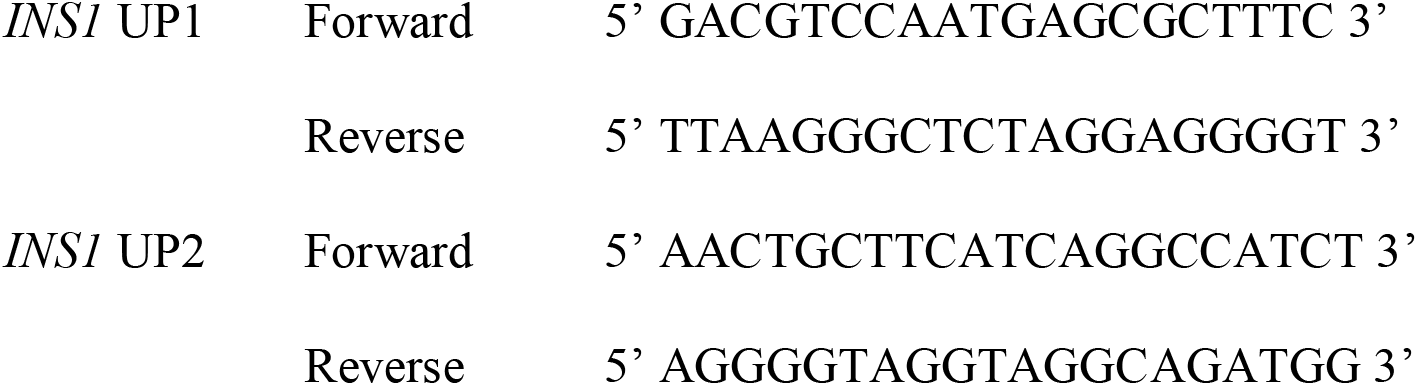
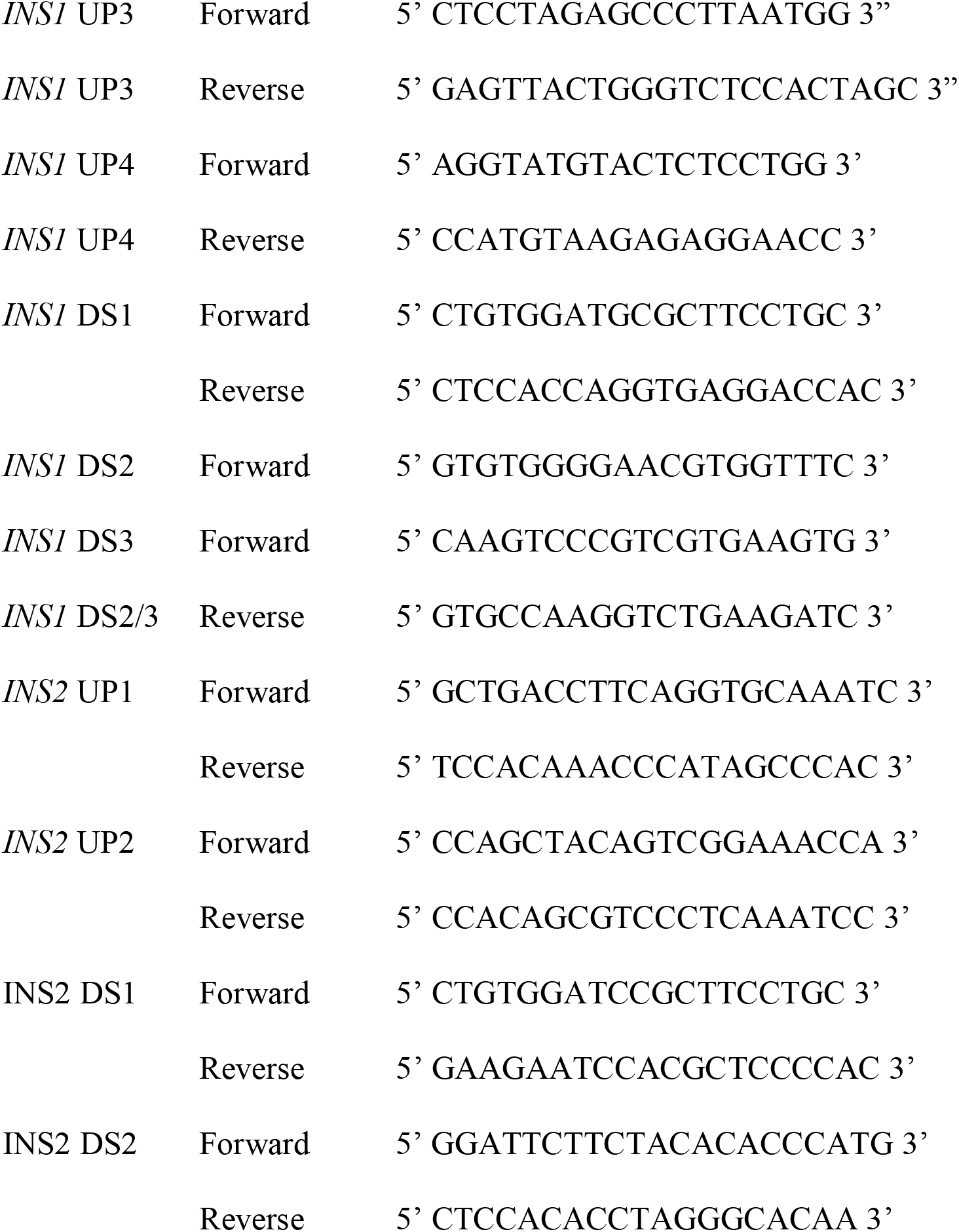

### Immunocytochemistry

Cells were plated at 50% confluency in poly-D-lysine coated 4-chamber glass bottom dishes (Cellvis) 16-24 hours prior to fixation. The following day cells were washed once with PBS, then fixed in 4% paraformaldehyde in PBS for 10 min at room temperature (RT). Cells were then washed three times with PBS and permeabilized in 0.2% Triton X-100 in PBS for 10 min at RT. Cells were blocked in 3% BSA in PBS for 1 hour at RT then incubated in primary antibody (mouse anti-AHCY 1:200 or mouse anti-PIP_2_ 1:50) overnight at 4°C. Following overnight incubation cells, were washed then incubated with anti-mouse IgG-κ Fc binding protein CFL 488 diluted 1:2000 in 3% BSA for 1 hour at RT. After 3 washes, cells were incubated with 5 µg/mL Hoechst 33342 in PBS for 10 min at RT. Hoechst 33342 solution was then removed after 10 min and cells were imaged in PBS by confocal microscopy on a Nikon A1Rsi confocal microscope.

### pERK1/2 assay-pERK1/2 assay

Cells were plated at 70% confluency in 24-well plates and incubated overnight at 37°C. 16-24 hours prior to assay, cells were incubated in serum-free low glucose RPMI-1640 media supplemented with 0.1% fatty acid-free BSA (FAF-BSA) overnight at 37°C, 5% CO_2_. Cells were washed once with PBS and pre-incubated with 200 µL KRBH containing 2.5 mM glucose for 30 minutes at 37°C, 5% CO2. After 30 minutes, KRBH was removed and replaced with either 200 µL KRBH containing the indicated concentrations of stimulants and were stimulated for 10 minutes at 37°C, 5% CO2. Supernatants were discarded and cells were lysed 75 µL TBS containing 1% Triton-X100 supplemented with protease and phosphatase inhibitors (20 mM sodium fluoride, 2 mM sodium orthovanadate, 10 mM β-glycerolphosphate, and 10 mM sodium pyrophosphate). 50 μg of each lysate was separated by SDS-PAGE on 10% acrylamide gels at 150 V for 90 min. Proteins were transferred onto PVDF membranes in ice-cold Towbin Buffer, 10% ethanol at 100V for 1 hour. Membranes were blocked in 3% BSA in TBST for 1 hour at room temperature and then incubated with anti-ERK (1:2000) and anti-pERK (1:1000) in 3% BSA in TBST overnight at 4°C. Primary antibody was removed the following day, and membranes were incubated in goat anti-mouse IgG HRP (1:10,000) and goat anti-rabbit IR800 (1:10,000) in 3% BSA in TBST for 1 hour at room temperature. Chemiluminescence was detected using standard ECL reagents, and chemiluminescence and fluorescence were imaged on Azure Biosystems Sapphire imager.

### LC-MS/MS analysis of proteins

Cells were lysed using a Barocycler (5°C, 60 cycles: 50 s at 35,000 psi and 10s at 1 atmospheric pressure) in 100 mM ammonium bicarbonate. Protein concentration was measured by bicinchoninic acid (BCA) assay (Pierce) and 50 ug of total protein for each sample was precipitated with 4 volumes of cold acetone (-20°C), and used for sample preparation as described previously [54, 55]. Dried and C18-cleaned peptides were re- suspended in 96.9% purified water, 3% acetonitrile, and 0.1% formic acid at a 1μg/μL, and 1µL was used for LC-MS/MS analysis in the Orbitrap Fusion Lumos mass spectrometer (Thermo Fisher Scientific) [54, 56]. The LC-MS/MS raw data were processed using MaxQuant (v1.6.3.3) [57] for protein identification and label-free quantitation [58]. MaxQuant results files were merged by matching rows based on the gene names. Proteins marked as “contaminants”, “reverse” and “only identified by match between runs” were removed. All LFQ values were then Log_2_ transformed for normalization, and samples were grouped based on deletions (control, IRBIT^KO^ or RYR2^KO^). Proteins identified in two replicates in at least one group (control or IRBIT^KO^; control or RYR2^KO^) were filtered for subsequent processing. Missing values were imputed using a constant (Zero-fill), and the average Log_2_(LFQ) values for each group were then calculated. Downregulated proteins were considered as proteins with Log_2_(Fold-Change) < -1 and average MS/MS count ratio < 0.5, compared to control. Similarly, upregulated proteins were considered as proteins with Log_2_(Fold-Change) > 1 and average MS/MS count ratio > 2, compared to control.

### Gene Ontology (GO) Analysis

Proteins identified as up- or down-regulated in RyR2^KO^ or IRBIT^KO^ cells were analyzed for over-representation in specific cellular component, biological process, or molecular function categories using PANTHER version 16 (http://www.pantherdb.org) [59]. Results were filtered at FDR < 0.05, > 2-fold enrichment, and a minimum of 4 proteins per category, except for IRBIT^KO^ down-regulated proteins which were filtered at FDR < 0.05 only.

## Acknowledgments

The authors thank Dr. Wayne Chen, Department of Physiology and Pharmacology, Libin Cardiovascular Institute, Calgary, Alberta Canada for the gift of a plasmid containing mouse RyR2 cDNA fused to GFP cDNA. We thank Drs. Robert Stahelin, Benita Sjogren, and Darcy Trader, Medicinal Chemistry and Molecular Pharmacology Department, Purdue University, for gifts of various reagents, and advice regarding their use. We thank Dr. Justin LaVigne, Medicinal Chemistry and Molecular Pharmacology Department, Purdue University, for advice regarding rt-qPCR experiments. All LC-MS/MS analysis including sample preparation, data collection and analysis were performed at the Purdue Proteomics Facility, Bindley Bioscience Center. The authors thank Rodrigo Mohallem for help with heatmaps.

## Funding

This work was supported by a Richard and Anne Borch Award (G.H.H.) and Showalter Faculty Scholar Award (G.H.H.).

## Author Contributions

K.E.H. and E.K.L. conducted most of the experiments and performed data analysis. P.A.S. contributed to rt-qPCR experiment. E.P.S.P and A.E.S contributed to the construction of knockout cell lines. M.S.D performed the methylation-dependent PCR experiments. U.A. conducted the LC-MS/MS experiments and data analysis. H.G. contributed to the design and analysis of the methylation-dependent PCR experiments. All authors contributed to the writing of the paper. G.H.H. designed the study, analyzed data, and wrote the paper.

## Competing interests

The authors declare no competing interests.

## Data and materials availability

All data needed to evaluate the conclusions in the paper are present in the paper or the supplementary materials. The LC-MS/MS raw data are deposited in MassIVE (massive.ucsd.edu), a publicly accessible data repository, with ID MSV000088343.

## Supplementary Materials

Fig. S1 Proteins with increased abundance in RyR2^KO^ cells

Fig. S2 Proteins with decreased abundance in RyR2^KO^ cells

Fig. S3 Proteins with increased abundance in IRBIT^KO^ cells

Fig. S4 Proteins with decreased abundance in IRBIT^KO^ cells

## Notes

https://massive.ucsd.edu

### Competing Interest Statement

The authors have declared no competing interest.

