## Supplementary material for "RyR2/IRBIT regulates insulin gene transcription, insulin content, and secretion in the insulinoma cell line INS-1": Supplmental Figures 1, 2, 3, & 4

**Supplementary Materials**

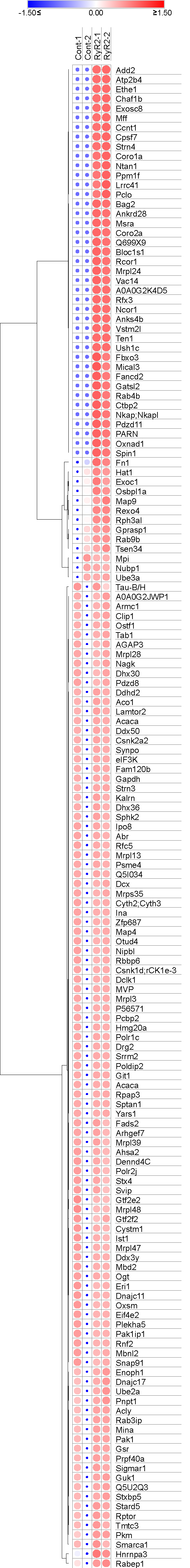

**Fig. S1. Proteins with increased abundance in RyR2^KO^ cells compared to control INS-1 cells as determined by LC-MS/MS**

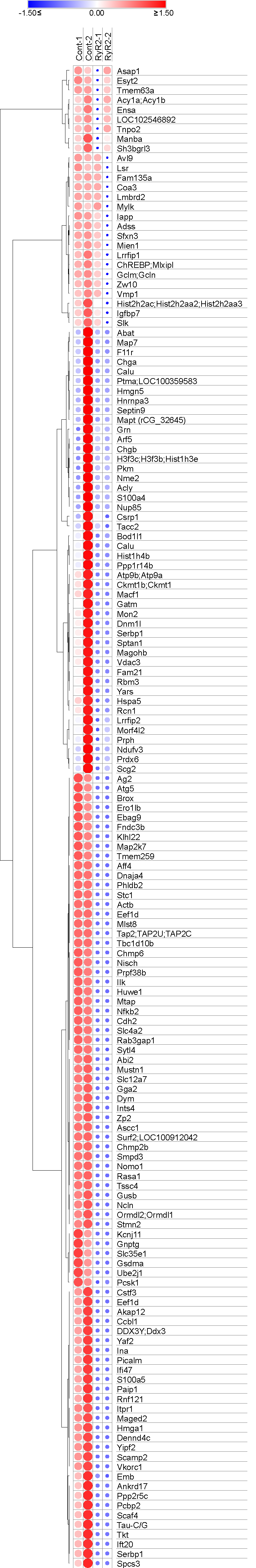

**Fig. S2. Proteins with decreased abundance in RyR2^KO^ cells compared to control INS-1 cells as determined by LC-MS/MS**

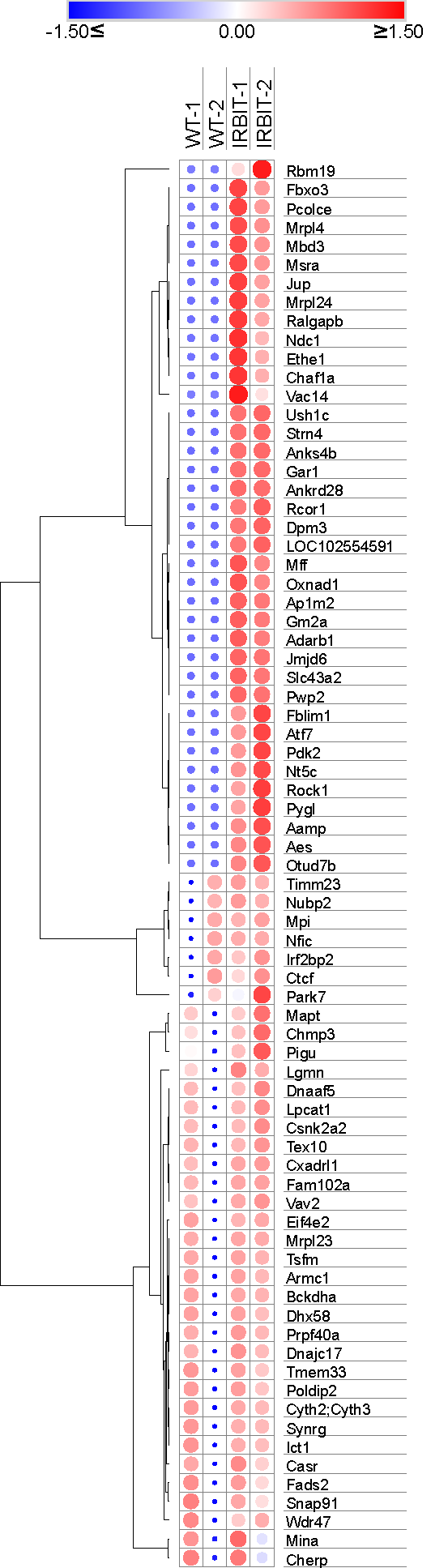

**Fig. S3. Proteins with increased abundance in IRBIT^KO^ cells compared to control INS-1 cells as determined by LC-MS/MS.**

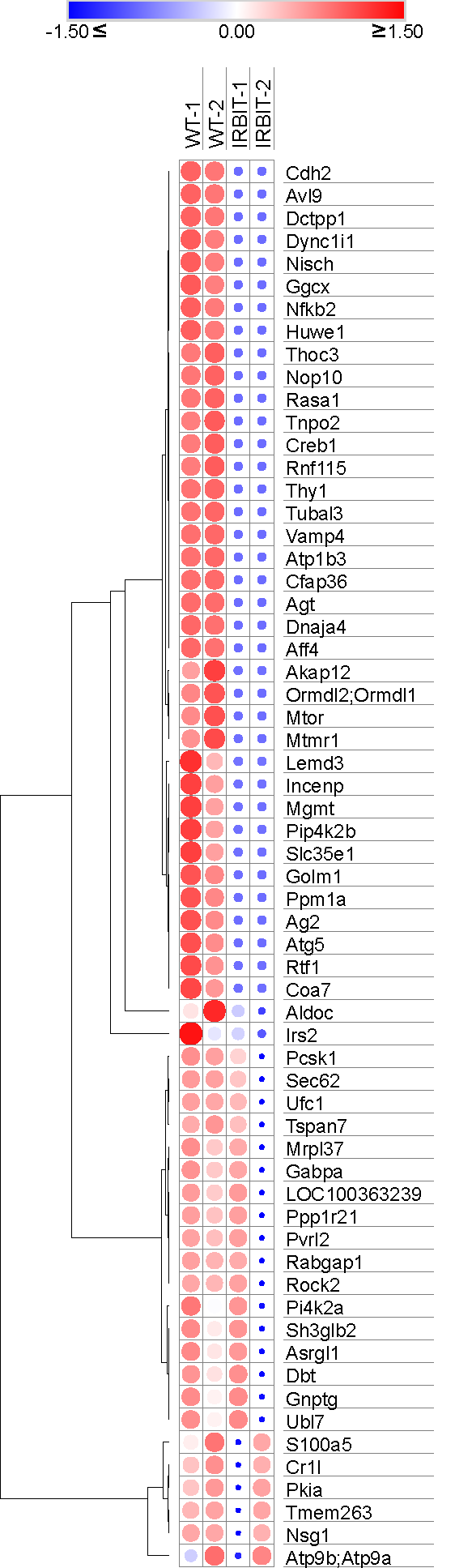

**Fig. S4. Proteins with reduced abundance in IRBIT^KO^ cells compared to control INS-1 cells as determined by LC-MS/MS.**
